# A distinctive tumor compartment in pancreatic lobules defined by nascent stroma and classical tumor cell phenotype

**DOI:** 10.1101/2024.03.14.584614

**Authors:** Sara Söderqvist, Annika Viljamaa, Natalie Geyer, Carina Strell, Neda Hekmati, Jennie Engstrand, Ernesto Sparrelid, Caroline Salmén, Rainer L. Heuchel, Argyro Zacharouli, Poya Ghorbani, Sara Harrizi, Yousra Hamidi, Olga Khorosjutina, Stefina Milanova, Bernhard Schmierer, Béla Bozóky, Carlos Fernández Moro, Marco Gerling

## Abstract

Pancreatic ductal adenocarcinoma (PDAC) is a highly aggressive tumor type characterized by a particularly extensive stroma. While different types of cancer-associated fibroblasts (CAFs) in this desmoplastic stroma have been described, areas of early invasion and nascent stroma are understudied. Here, we identify a distinctive PDAC niche within the pancreatic lobules, a compartment dominated by pancreatic exocrine cells and slender stroma. Cellular interaction profiling using machine learning on whole slide images of human PDAC reveals that the tumor invasion front in the lobules is dominated by specific interactions of tumor cells and exocrine cells that have undergone acinar-to-ductal metaplasia (ADM). Multiplex protein and mRNA stains confirm that tumor growth in the lobules is closely linked to ADM in the lobules, and reveal stromal protein gradients from the gracile lobular stroma to the characteristic desmoplastic stroma. We identify nascent CAFs (nCAFs), co-expressing expressing nerve growth factor receptor (NGFR) and platelet-derived growth factor receptor alpha (PDGFRa) that are absent in the mature, desmoplastic stroma. Lobular invasion and nCAFs are intertwined with phenotypic changes of the cancer cells, such that tumor cells in lobules express classical subtype markers, while those embedded in the desmoplastic are on the basal end of the phenotypic continuum. In mice, the PDAC subtype – basal or classical – similarly depends on tissue location, suggesting microenvironmental factors rather than clonal selection as important drivers of tumor phenotype identity. Clinically, our results mandate factoring in tumor tissue location when calling PDAC subtypes. Biologically, they identify pancreatic lobules as a distinctive tissue niche associated with nascent stroma, and they suggest that lobular colonization by tumor cells is a significant route of PDAC progression.

## Introduction

Pancreatic ductal adenocarcinoma (PDAC) is the main histological type of pancreatic cancer and has one of the worst survival rates among all solid tumors^1^. PDAC histology is dominated by an extensive desmoplastic stromal reaction that surrounds tumor cell nests^2^. This desmoplastic microenvironment is thought to result from stromal cell proliferation and extracellular deposition of matrix proteins^2^, largely driven by tumor cell-derived signals^3–7^. An extensive body of research has revealed heterogeneous stromal populations around PDAC tumor nests with both tumor-promoting and -restraining functions^2^, such as myofibroblastic cancer-associated fibroblasts (myCAFs) and inflammatory CAFs (iCAFs), which have partially opposing roles in PDAC progression^8,9^.

In contrast to PDAC, the initial stages of its most common precursor lesion, pancreatic intraepithelial neoplasia (PanIN), lack a desmoplastic stroma, which starts to develop first in more advanced PanIN stages^10^. Mouse models have revealed that oncogenic transformation of pancreatic acinar or duct cells precedes desmoplasia formation by days to weeks^11,12^. Together, these data establish a sequence where oncogenic transformation of epithelial cells leads to stromal activation. In established tumors, where cancer cells invade healthy tissues, the corresponding sequence of events is largely unknown. Particularly, areas of nascent desmoplasia have not been systematically defined.

Two main molecular PDAC subtypes have been identified based on RNA sequencing, “basal” (or “basal-like”) and “classical”^13–16^. Basal tumors have dismal outcomes and are largely resistant to chemotherapy^15^. In individual tumors, basal and classical cells as well as their hybrid forms co-exist^17,18^. Recent spatially resolved data have started to connect tumor subtypes to distinct stromal cells^18^, suggesting that stromal factors, at least partially, drive tumor cell phenotypes. These observations imply significant microenvironmental control of tumor cell behavior, and they suggest that calling tumor subtypes needs to take the tissue context into consideration, which has significant implications on the use of the subtypes in clinical practice and when using them for stratification in clinical trials.

Owing to the lack of recognition of early invasion in PDAC, most studies associating tumor and stromal phenotypes have focused on the different stroma-rich compartments and disregarded regions of early-activated, ‘nascent’ stroma^18,19^. Hence, tumor cell phenotypes in areas of early tumor invasion into the non-malignant pancreatic parenchyma are insufficiently described.

Here, we use multiplex-immunohistochemistry and immunofluorescence on resected human PDAC to show that cancer cells colonize and invade the pancreatic lobules, where an area of nascent desmoplasia formation is located. Machine learning (ML)-based image analysis reveals that during lobular invasion, PDAC cells interact closely with a subset of non-malignant epithelial cells that have undergone acinar-to-ductal metaplasia (ADM). Quantitation of tumor phenotypes and stromal markers associates PDAC cells in the pancreatic lobules with the classical subtype and reveals concomitant stromal evolution, marked by the loss of nerve growth factor receptor (NGFR) expression, in progressing tumor colonization. In the stroma-rich, desmoplastic regions, PDAC cells adopt a basal-like phenotype close to tumor-promoting myCAFs. In mice, orthotopic injection of near-clonal tumor cells is associated with a switch to classical phenotypes in areas of acinar invasion. Together, our data connect tumor tissue location with tumor cell phenotypes, and they suggest a model of PDAC progression in which colonization of pancreatic lobules is a significant mode of tumor invasion.

## Results

### Invasion into pancreatic exocrine lobules defines a distinct compartment in PDAC

In histopathological assessment of resected PDAC, tumor cells are mainly seen embedded in a characteristic desmoplastic stroma (**Figure 1a**). However, we frequently observed in clinical routine that PDAC cells also invade the pancreatic lobules (**Figure 1a**), the major anatomic units of the pancreas that comprise acinar cells, the endocrine islets of Langerhans, intralobular ducts and slender connective tissue as their main components^20^. Because the peritumoral pancreas frequently shows chronic pancreatitis and reactive changes (**Figure 1a**), which impair the accurate identification of tumor cells^21^, we employed immunohistochemistry (IHC) for p53 to distinguish tumor cells from non-malignant lobular cells. In ***TP53*** mutated tumors, p53 protein accumulates, and its abundance in the nucleus revealed extensive tumor colonization of partially degraded lobules (**Figure 1b&c**). In another subset of tumors, SMAD family member 4 (SMAD4) protein expression is lost upon ***SMAD4*** mutation (**Figure 1d**), together with p53 allowing us to assess lobular invasion systematically in most cases. We reviewed the hematoxylin & eosin (H&E), and the corresponding IHC stains of *n* = 31 individual PDACs (**Table 1**). We found tumor cells in lobules in all cases analyzed (100%), although at varying extents and degrees of lobular invasion and atrophy (**Supplementary Figure 1a-c**). We quantified the fraction of tumor cells in lobules (tumor^in_lobules^) and desmoplastic stroma (tumor^in_stroma^) in whole-slide images (WSIs) of H&E-stained slides for a subset of *n* = 5 tumors, selected to represent different degrees of lobular invasion. We defined tumor^in_stroma^ as areas devoid of any lobular remnants, such as endocrine islets of Langerhans, which regularly remain discernable even in end-stage lobular atrophy (**Supplementary Figure 1d&e**). The proportion of lobular invasion of the total tumor area ranged from 4% to 32% (mean: 14.8%, standard deviation: 10.8%, **Figure 1e**). As expected, tumor^in_stroma^ – the histopathological archetype of PDAC – was the predominant tumor location; however, tumor^in_lobules^ emerged as a prevalent compartment of PDAC invasion.

**Figure 1.**
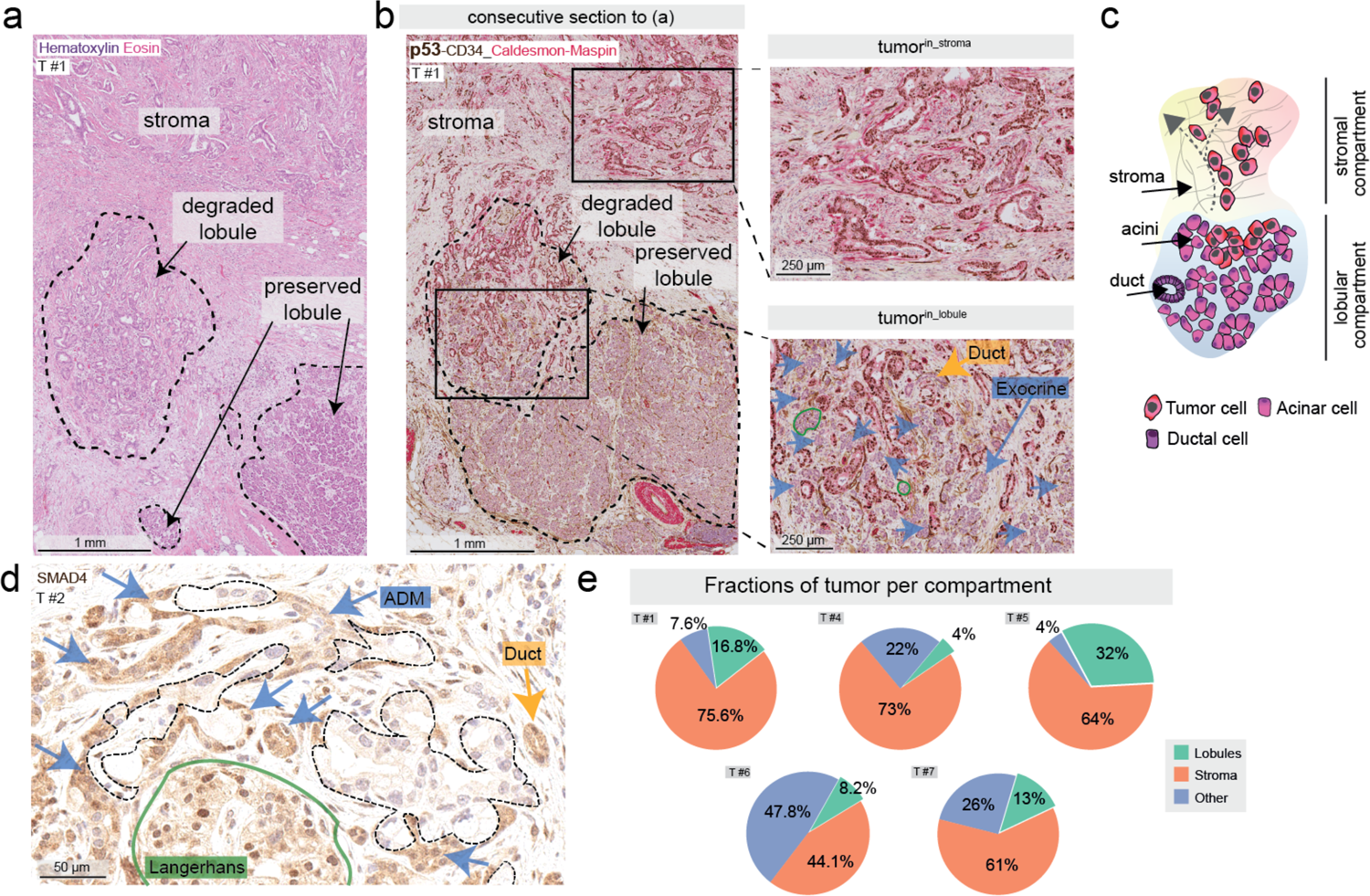
Pancreatic ductal adenocarcinoma growth inside the pancreatic lobules. **a)** Hematoxylin & eosin stain of a pancreatic ductal adenocarcinoma in desmoplastic stroma (upper part of the image, “stroma”), and pancreatic exocrine lobules (lower part of the image, dashed line) **b)** Immunohistochemistry (IHC) stain for the indicated proteins, including p53 (brown) of a consecutive section corresponding to (a). Dashed lines: pancreatic lobules. Green filled lines: endocrine islets of Langerhans, blue arrows: exocrine cells, yellow arrows: duct. **c)** Depicted schematically, tumor cells localize in the stroma-transformed compartment, or in the lobular structures of the pancreas. **d)** Annotated tumor nests with loss of SMAD family member 4 (SMAD4, brown, nuclear and cytoplasmic) in proximity to exocrine, ductal and endocrine cells. **e)** Annotations of all tumor comprising one whole-slide image from 5 individual tumors were classified to stromal invasion, acinar invasion and other. While the largest fraction of tumor is located in the stroma (44.1 – 75.6%), all tumors include a fraction residing in the lobule (4.8 - 32%, green). “Other” invasion represents tumor present in, for example, larger ducts (bile or pancreatic), adipose tissue and nerve, accounting for 4 - 47.8%. **a), b), d)** Representative of *n* = 31 tumors from *n* = 31 patients.

**Table 1:** Basic characteristics for the patients whose tumors were analyzed with multiplex immunohistochemistry. n/a: not available. TNM classification according to the pathological statement. Resection margin status was determined using the presence of tumor cells < 1 mm from any resection margin. Number of stains quantified for individual tumors are given in the Supplementary Table 1.

|  | *variable* | *value* |
| --- | --- | --- |
| **Age** | Median, years (min-max) | 70 (45-84) |
| **Sex** | male | 14 |
|  | female | 17 |
| **Tumor stage** | T1 | 0 |
|  | T2 | 16 |
|  | T3 | 14 |
|  | T4 | 1 |
| **Nodal stage** | N0 | 3 |
|  | N1 | 12 |
|  | N2 | 16 |
| **Metastatic stage** | M0 | 22 |
|  | M1 | 7 |
|  | n/a | 2 |
| **Lymphatic vessel infiltration** | pL0 | 0 |
|  | pL1 | 31 |
| **Venous infiltration** | pV0 | 4 |
|  | pV1 | 26 |
|  | n/a | 1 |
| **Perineural infiltration** | pPn0 | 2 |
|  | pPn1 | 29 |
| **Resection margin** | R0 | 10 |
|  | R1 | 21 |
| **Tumor Location** | Caput | 25 |
|  | Collum | 1 |
|  | Cauda | 5 |
| **Neoadjuvant chemotherapy** | Yes | 5 |
|  | No | 26 |
| **Relapse** | Yes | 25 |
|  | No | 6 |
| **Time-to-relapse** | Median, months (min-max) | 14 (2-46) |

### The classical PDAC cell subtype dominates in pancreatic lobules

The prognostic PDAC subtypes, classical and basal, are subject to microenvironmental control, which at least partly explains intratumoral subtype heterogeneity^22^. In the lobular microenvironment, tumor cells are not surrounded by a desmoplastic stroma. Instead, they establish direct contact with the adjacent non-malignant epithelial cells, reminiscent of the replacement-like pattern of tumor growth described in other organs^23^. Hence, tumor cells in lobular areas are likely exposed to different paracrine signals than tumor cells in the stroma^18,22^.

Independent studies have shown that a limited number of markers can approximate subtype identity on a single-cell level in situ^24,25^. Here, we used multiplex IHC with a panel of six markers, two classical (caudal type homeobox 2 [CDX2], mucin 5 subtypes A and C [MUC5AC]), and four basal markers (Cytokeratin (CK)5, high mobility group AT-hook 2 [HMGA2], Carbohydrate antigen (CA)125/MUC16, and CK17) to approximate subtype identity of tumor^in_lobules^ and tumor^in_stroma^ (**Figure 2a**). We annotated tumor cell regions on WSIs and built classifiers in QuPath^26^ to quantify positive tumor cells in each tissue location (**Figure 2b&c**, **Supplementary Figure 2a**). As expected, there was a strong positive correlation between the classical markers, MUC5AC and CDX2, and between the basal markers, CK5, HMGA2, CA125, and CK17, respectively (**Supplementary Figure 2b**).

**Figure 2.**
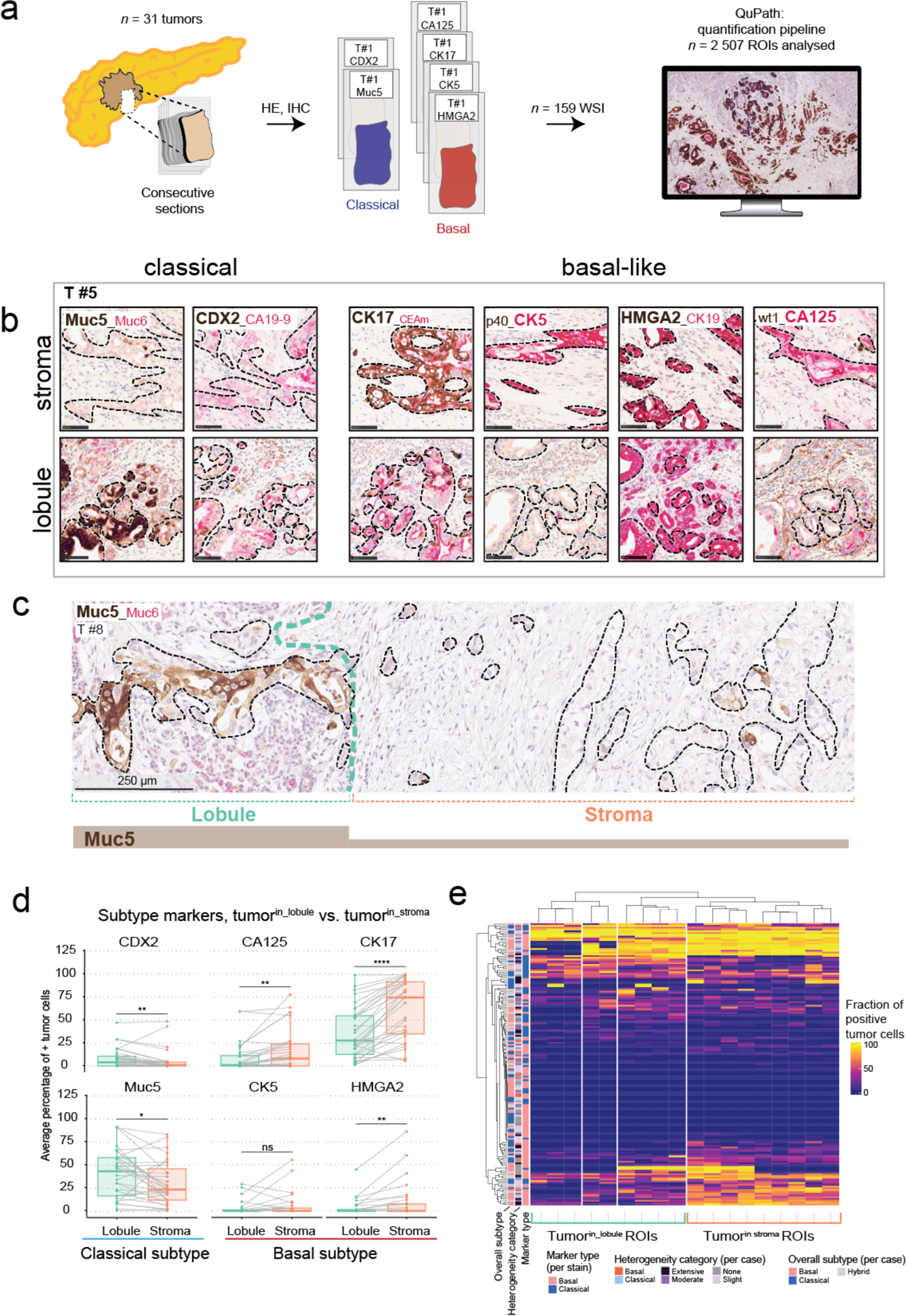
Subtype-related expression depends on the tissue compartment. **a)** Schematic of the sampling strategy to analyze compartment-dependent heterogeneity in pancreatic ductal adenocarcinoma. Consecutive sections of *n* = 31 tumors were stained with various Immunohistochemistry (IHC) combinations as part of the diagnostic workflow at Karolinska University Hospital. Out of these, *n* = 2 classical-related stains and *n* = 4 basal-related stains were analyzed in QuPath**. b)** Representative IHC stains of subtype-related expression patterns of tumor cells residing in the stroma (upper panel) or lobule (bottom panel) of one tumor. From left to right: The classical stains, mucin 5 subtypes A and C (MUC5AC, brown, cytoplasmic) and caudal type homeobox 2 (CDX2, brown, nuclear) and the basal-like related stains, Cytokeratin (CK)17 (brown, cytoplasmic), CK5 (red, cytoplasmic), high mobility group AT-hook 2 (HMGA2, brown, nuclear), Carbohydrate antigen (CA)125 (red, cytoplasmic). Scale bars: 100 µm. Representative of *n* = 31 tumors. The stromal and acinar regions of interest are overlapping regions of consecutive sections. **c)** Representative IHC of MUC5AC where lobular invasion (left) is denoted by expression of the classical marker, which is lost at the stromal invasion (right). Black dashed line: tumor, turquoise dashed line: lobule. **d)** paired Wilcoxon rank sum test of differential expression in stromal or lobular tumor locations. Boxplots show quartiles of the means from all regions of interest (ROIs) dichotomized to stromal - or lobular invasion of *n* = 31 tumors with Benjamini-Hochberg correction for multiple testing. **e)** Unsupervised clustering of the fractions of positive tumor cells for each tumor and stain combination over all quantified ROIs (*n* = 2507). The metadata categories describe marker type, as classical and basal. Heterogeneity category represents the number of stains that were significantly differentially expressed for each tumor; None; no stain, Slight; one stain, Moderate; two-three stains and Extensive; 3 or more stains. Overall heterogeneity was assessed by the fraction of all positive cells for the subtype-related stains. ns: non-significant, * = 0.05, ** = 0.01, 0.01, **** = 0.0001. **b**, **c**: Representative of *n* = 31 tumors.

When we compared classical and basal marker expression between tumor^in_lobules^ and tumor^in_stroma^, we found significantly lower fractions of cells positive for basal markers CA125, CK17, and HMGA2. In contrast, the fractions of tumor cells positive for the classical markers, MUC5AC and CDX2 were higher in tumor^in_lobules^ as compared to tumor^in_stroma^ (**Figure 2d and Supplementary Figures 3 - 8**). Accordingly, unsupervised hierarchical clustering of stain quantitations separated the regions of interest (ROIs) in the lobular compartment from stromal invasion (**Figure 2e**).

To assess the overall degree of heterogeneity in all quantified regions irrespective of lobular or stromal location, we used the Simpson diversity index, an established heterogeneity metric^27,28^. Here, the Simpson index describes the probability of retrieving two random ROIs with different fractions of positive tumor cells of each respective subtype marker, allowing us to assess overall heterogeneity per tumor and stain. We found a poor correlation of the Simpson index with compartment-dependent differential expression between tumor^in_lobules^ and tumor^in_stroma^ (**Supplementary Figure 9**), suggesting that tumor cell location in lobules *vs*. stroma is one of several determinants of subtype identity.

Our data show that the pancreatic lobules represent a distinctive tumor compartment associated with the predominance of classical-type tumor cells, implying microenvironmental control of subtype identity.

### Machine learning-driven interaction mapping identifies cells undergoing acinar-to-ductal as main tumor interaction partners

Tumor^in_stroma^ represents the archetype histological pattern of PDAC, in which tumor cells are embedded in and physically connect with stromal cells and the extracellular matrix (ECM)^20^. Tumor cells in lobules meet different cellular interaction partners, potentially comprising all the cell types of the pancreatic lobules. To quantitate cellular interactions in tumor^in_lobules^ *vs*. tumor^in_stroma^, we trained a convolutional neural network to recognize the main tissue types on WSIs stained with IHC for NGFR and cluster of differentiation (CD)146. Compared to H&E stains, NGFR/CD146 improves contrast to differentiate, for example, stromal and epithelial structures. We trained the model to recognize acinar cells, exocrine cells that had undergone ADM, ductal cells, immune infiltrates, fibroblasts, and ECM deposits (summarized as ‘fibrosis’), vessels, nerves, endocrine islets of Langerhans, and tumor cells, based on the main distinguishable tissue classes (**Figure 3a, Supplementary Figure 10a**). The resulting ML-based annotations (*n* = 9326 annotations from *n* = 31 ROIs in *n* = 5 tumors), reliably identified tumor cells and all other tissue types with good accuracy, allowing us to assess the contacts of tumor cells to other tissue types in situ (**Figure 3b**). After benchmarking against manual annotations (**Supplementary Figure 10b**), we quantitatively compared tumor cell contacts to all tissue types in tumor^in_lobules^ (*n* = 15 ROIs) with tumor^in_stroma^ (*n* = 26 ROIs). We found that PDAC cells in both compartments were most frequently adjacent to ‘fibrosis’ (**Figure 3c**). However, tumor^in_lobules^ interacted less frequently with the ‘fibrosis’ (97% in tumor^in_stroma^ vs. 80.5% in tumor^in_lobules^), and established exclusive interactions with ‘acinar’ and ‘ADM’ classes, which were absent in tumor^in_stroma^, as expected (**Figure 3c**). In the stroma-poor areas of the tumor-acinar interface in the lobules, tumor cells predominantly interacted with acinar cells that had undergone ADM (**Figure 3d**). Multiplex RNA in situ hybridization (ISH) confirmed the upregulation of both the confirmed ADM markers, C-reactive protein (***CRP***), regenerating family member (***REG****)**3A**,* and the proposed ADM marker melanoma cell adhesion molecule (***MCAM****)/**CD146***, such that double positive cells for the acinar marker, carboxypeptidase (***CP****)**B1***, and the different ADM markers were seen in the invaded acini, although at different distances to the invading Keratin (***KRT****)**19**^+^*tumor cells (**Figure 3e**)^18,29^.

**Figure 3.**
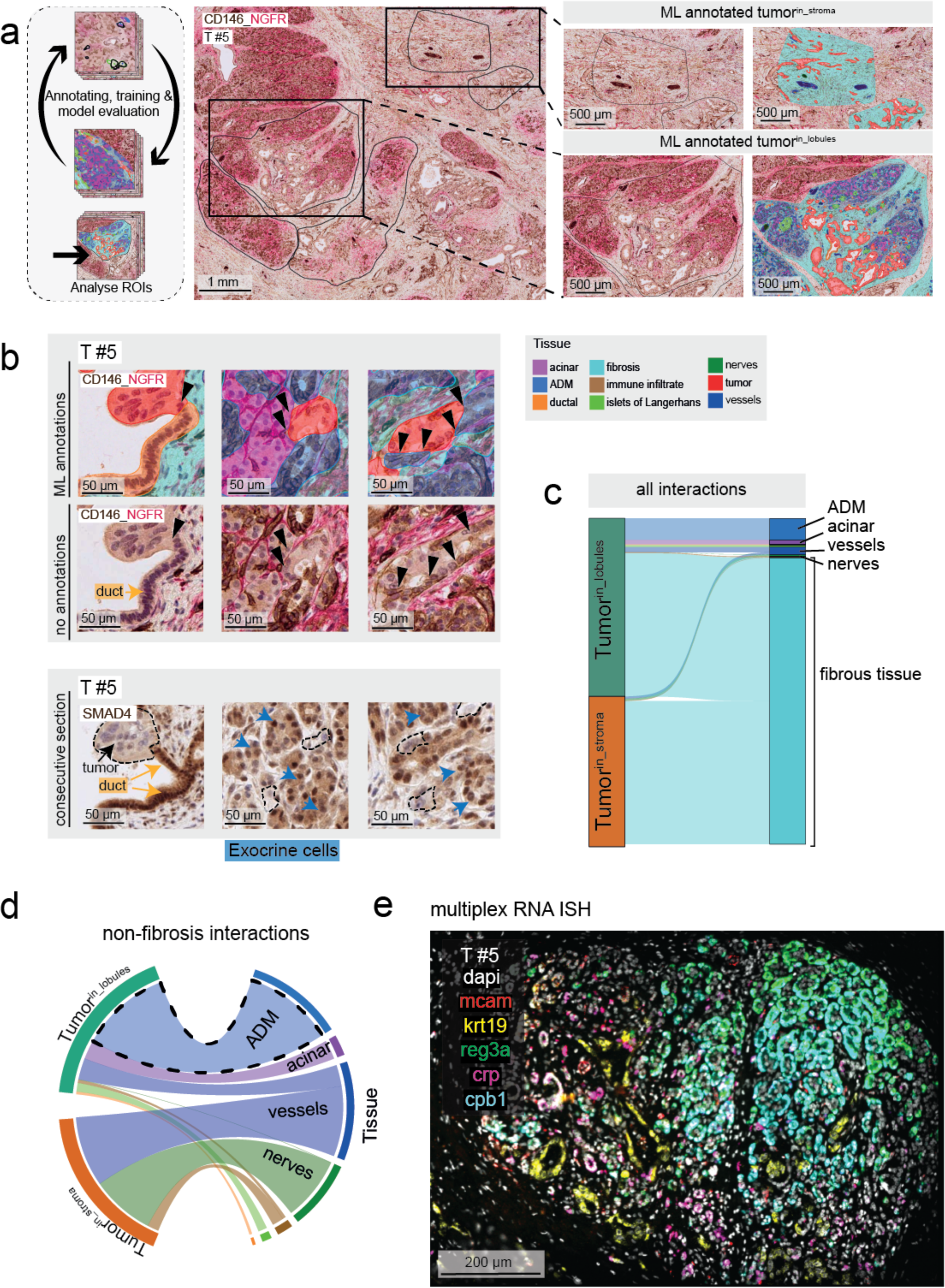
Tumor cells in lobules interact with unique cell types at the lobular invasion front. **a)** A convolutional neural network model was trained to recognize the main cellular and tissue architecture types in pancreatic ductal adenocarcinoma (left panel). Detection of multiple classes in stroma (upper panel) and lobule (lower panel) are shown, where the machine learning derived annotations are overlayed on the right inserts. Representative of 5 tumors. **b)** The tumor (red) interactions with the nearest class from left to right (upper panel): ‘duct’ (orange overlay), ‘acinar’ (magenta overlay) and acinar-to-ductal metaplasia (‘ADM’, dodger blue overlay). Tumor nests with loss of SMAD family member 4, ([SMAD4], brown, nuclear and cytoplasmic) on a consecutive section (lower panel). The black arrows indicate individual interacting cells; blue arrows point at exocrine cells and yellow arrows mark ducts. Representative of *n* = 5 tumors. **c)** Quantitative comparison of tumor cell contacts to all tissue types. Data from *n* = 5 tumors; *n* = 15 lobular regions of interest (ROIs) (tumor^in_lobules^) and *n* = 26 stromal ROIs (tumor^in_stroma^). **d)** Quantitative comparison of tumor cell contacts to all tissue types, besides ‘fibrosis’ class. Data from *n* = 5 tumors; *n* = 15 lobular ROIs (tumor^in_lobules^) and *n* = 26 stromal ROIs. **e)** multiplex-RNA in situ hybridization of lobular invasion. Tumor and other ductal structures (Keratin (***KRT****)**19**^+^*, yellow) surrounded by acinar cells (Carboxypeptidase *[**CP**]**B1**^+^*, cyan) co-expressing ADM-markers (Melanoma cell adhesion molecule *[**MCAM**]*, red; C-reactive protein *[**CRP**]*, magenta; and Regenerating family member [***REG****]**3A***, green) in different combinations. Representative image of *n* = 2 tumors.

Hence, invasion into the lobular compartment is strongly associated with the presence of ADM, where acinar cells that have undergone ADM are the primary interaction partners of cancer cells, defining the leading edge of PDAC tumor invasion in the lobules.

### A distinct type of NGFR+/PDGFRa+ fibroblasts dominates in the invaded lobules

Despite prevalent contacts of tumor^in_lobules^ with the exocrine epithelium, fibrosis-tumor contacts were also frequent in the lobules (**Figure 3c**). Using the ML-annotated data to generate spatial maps of tumor-tissue interactions (**Supplementary Figure 11)**, we found that in the lobules, interactions of PDAC cells and the ‘fibrosis’ class were enriched at the periphery of the lobules, while interactions between tumor cells and ADM cells peaked at the leading edge of the tumor-exocrine cell border (**Figure 4a**). These data suggested that stromal activation and desmoplasia increase with time after tumor cell colonization of the lobules.

**Figure 4.**
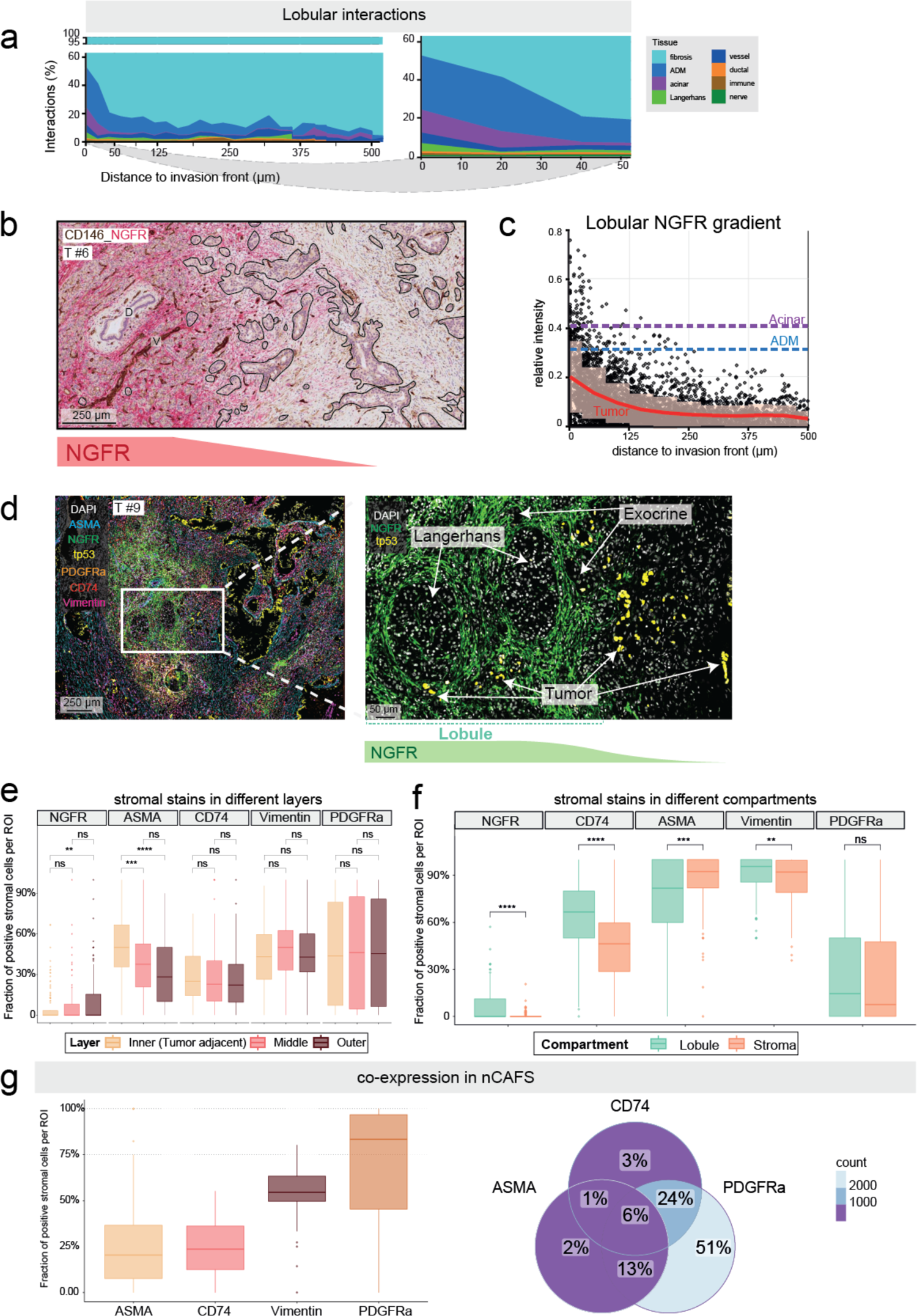
Distinct stromal cells in lobular and stromal compartments. **a)** Quantification of the distances of tumor cell interactions from the lobular invasion front. Right panel: magnification of the first 50 µm; combined data from *n* = 5 tumors, *n* = 15 lobular regions of interest (ROIs). **b)** Representative image of immunohistochemistry (IHC) stains for nerve growth factor receptor (NGFR), costained with cluster of differentiation (CD)146 (brown, cytoplasmic). Representative of 4 tumors. Black lines indicate tumor nests; D: duct, V: vessel. **c)** IHC-based quantification of NGFR expression of the area expanded from tumor cells (individual data points). Red line: rolling average of tumor cell-based measurements with standard deviation. Magenta and dodger blue lines; averages of NGFR expression based on the expanded area of acinar and acinar cells undergoing acinar-to-ductal metaplasia, respectively (*n* = 4 tumors, *n* = 12 lobular ROIs). **d)** Representative image of multiplex-immunofluorescence staining for the indicated proteins. Representative of *n* = 8 tumors. **e)** Differential distribution of stroma cells expressing the stromal proteins, NGFR, alpha-smooth muscle actin (ASMA), cluster of differentiation (CD)74, Vimentin and platelet-derived growth factor receptor alpha (PDGFRa) at three distinct distance layers from the tumor nests. Unpaired two-tailed Kruskal-Wallis rank sum test, with post-hoc Dunn’s test for pairwise multiple comparisons. **f)** Dichotomized stratification of stroma cell expression of the indicated proteins next to tumor^in_lobules^ and tumor^in_stroma^. Unpaired two-tailed Wilcoxon rank sum test, with Benjamini-Hochberg correction for multiple testing. **g**) Co-expression of protein markers in NGFR+ stroma cells in the lobules (nCAFs). Data from *n* = 8 individual tumors, with *n* = 53 ROIs in total. ns: non-significant, * = 0.05, ** = 0.01, *** = 0.001, **** = 0.0001. **e), f)**: data from *n* = 8 individual tumors, *n* = 363 ROIs (222 stromal and 141 lobular)

NGFR, a stromal marker that has been associated with favorable prognosis in PDAC and is upregulated in benign fibrotic conditions^30–33^, was strongly expressed in lobular regions (**Figure 4b**). To quantitate peritumoral NGFR expression in relation to progressing tumor cell invasion with increasing distance to the tumor-acinar border, we used the ML-generated annotations and measured NGFR intensity around acinar cells, cells showing ADM, and tumor cells (**Supplementary Figure 11)**. We found that NGFR expression was high in the proximity of acinar cells with and without signs of ADM; however, peritumoral NGFR expression decreased with increasing distance of the tumor cells to the intralobular invasion front (**Figure 4c**). These data identified NGFR^+^ cells as a distinct stromal cell population in regions of nascent PDAC invasion.

In desmoplastic tumor regions, previously established CAF identities can be approximated by alpha-smooth muscle actin (ASMA) for myCAFs, PDGFRa for iCAFs, and CD74 for antigen-presenting CAFs (apCAFs)^9^. To systematize NGFR^+^ stromal cells in the realm of known fibroblast markers, we used multiplex-immunofluorescence (m-IF) for ASMA, PDGFRa, CD74, the pan-stroma marker, Vimentin, and the tumor marker, p53, (*n* = 8 tumors, all of which were ***TP53*** mutated to be able to identify the tumor cells **Figure 4d & Supplementary Figure 12a**). We found ASMA^+^ cells, as expected^8^, to be significantly enriched in the stroma layer next to tumor cell nests in both stroma-transformed and lobular compartments, while PDGFRa^+^ cells and CD74^+^ cells, respectively, were enriched in neither of the three layers (**Figure 4e & Supplementary Figure 12b**). Tumor^in_stroma^ was deprived of NGFR (**Figure 4f & Supplementary Figure 12c)**. Based on their location next to invading PDAC cells in the lobules, we termed NGFR^+^ peritumoral cells “nascent CAFs” (nCAFs). nCAFs co-expressed PDGFRa to a high extent (94%), while 22% of nCAFs were positive for ASMA and 34% for CD74 (**Figure 4g & Supplementary Figure 13a**). In contrast, surrounding tumor^in_stroma^, PDGFRa+ cells were virtually negative for NGFR (**Supplementary Figure 13b-c**)

These data identify NGFR^+^/PDGFRa^+^ nCAFs in close proximity to tumor cells in early invasion of pancreatic lobules, and they show that NGFR is lost in the archetypical, desmoplastic regions.

### Compartment-driven phenotypic switch in clonal murine pancreatic cancer

Our data suggested interactions between tumor cell phenotypes and lobular *vs*. stromal tumor cell location, such that the classical phenotype is partly driven by tissue compartment. This niche effect could be due to the preference of individual tumor clones to colonize the lobules. Alternatively, lobular or stromal cues could induce the distinctive phenotypes. Distinguishing between both models is important for understanding PDAC evolution and invasion, because the underlying mechanism is essential for developing strategies to inhibit tumor invasion in either tissue compartment.

To test whether tumor and stromal phenotypes co-develop gradually or by clonal selection, we first analyzed tumors from a previous experiment^34^, derived from genetically engineered KPC mice (*LSL-Kras^G12D/+^;LSL-Trp53^R172H/+^;Pdx-1-Cre)*, removing some of the inherent heterogeneity of human tumors. KPC mice recapitulate human PDAC histopathology and develop a typical desmoplastic stromal reaction^35^. We used IHC for the basal marker, HMGA2, to assess tumor phenotypes, which revealed a decrease in HMGA2 abundancy in areas of lobular invasion (**Supplementary Figure 14**). These observations established the presence of lobular invasion in KPC mice and indicated environmental cues, rather than clonal selection, as drivers of tumor cell phenotypes. However, autochthonous KPC tumors have been shown to be multiclonal^36^. Hence, we could not rule out clonal selection as the driving factor of tumor cell phenotypes in this model.

To further reduce clonality as a factor for tumor diversity, we next used an orthotopic injection model, where KPCT cells (bearing an additional, inducible tdTomato allele, *B6.Cg-Gt(ROSA)26Sor^tm14(CAG-tdTomato)Hze^*/*J*)^37^ are injected orthotopically into fully immunocompetent C57BL/6 mice. First, we lentivirally integrated a library of barcodes into KPCT cells at low multiplicity of infection (**Figure 5a**). We injected 10e^5^ barcoded cells into the pancreata of *n* = 4 mice, collected the established tumors, and counted the barcodes by amplicon sequencing of the lentivirally integrated cassette. In three out of four mice, two clones comprised >90% of the tumor cells (**Figure 5b**). In two of those mice, tumors were monoclonal, such that at least 99% of the tumor cells expressed the same individual barcode, originating from one single cell (**Figure 5b**). Five clones comprised >90% of the tumor cells in one case. Hence, the injection of PDAC cells into the pancreas was associated with massive clonal selection, in line with recent data from another injection-based tumor model^38^.

**Figure 5.**
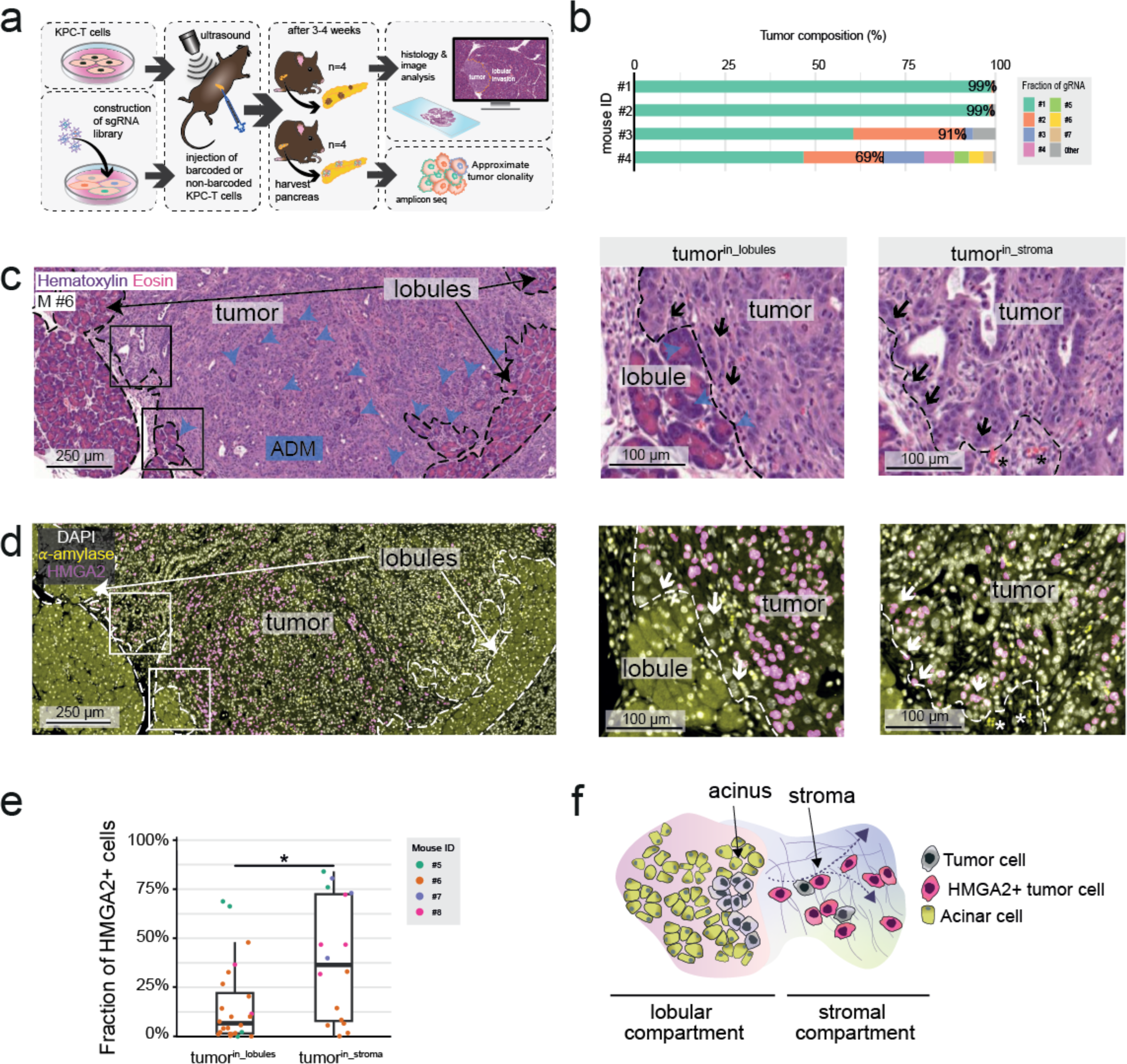
Compartment-dependent phenotypic switch in clonal tumors *in vivo*. **a)** Schematic of orthotopic injection model for analysis of subtype heterogeneity and clonality screen. KPCT cells were injected into the pancreas of C57BL/6J mice. 3-4 weeks after injection, mice were euthanized and the pancreata with tumor(s) harvested. A clonality screen was performed by constructing a sgRNA library of the KPC cells before injection, resulting in injection of 10e^5^ barcoded cells in four mice. **b)** Fractions of unique clones identified in the harvested tumors per mouse in the clonality screen. Data from *n* = 4 tumors. **c & d)** Representative (H&E) stain (**c**) and immunofluorescence of high mobility group AT-hook 2 HMGA2 (basal subtype marker, pink, nuclear) and α-amylase (yellow, cytoplasmic) on a consecutive section (**d**) of a mouse tumor from the orthotopic injection model (*n* = 4 tumors). Black or white arrows: tumor cells in direct contact with acini (middle panel) or stroma (right panel), blue arrows: acinar-to-ductal metaplasia (ADM), dashed line separates tumor from lobules and stroma. **e)** Fractions of HMGA2+ tumor cells of lobular invasion (tumor^in_lobules^) and stromal invasion (tumor^in_stroma^) from the orthotopic injection model. Data from *n* = 4 tumors and *n* = 40 regions of interest, the result from a two-sided Wilcoxon rank sum test with Benjamini-Hochberg multiple testing correction. * = 0.05. Box-and-Whisker plot, median (line), interquartile range (IQR; box), minimum and maximum values within 1.5 times the IQR from the first and third quartiles (whiskers) and individual data points are shown **f)** Illustration of the phenotypic switch as pancreatic ductal adenocarcinoma cells invade into two distinct tissue compartments with the basal-like phenotype dominating in stromal compartment (HMGA2+ tumor cells).

H&E stains revealed a stroma-rich tumor core upon orthotopic injection of KPCT cells, although the stroma was less dense than in the autochthonous model (**Figure 5c**), possibly owing to the limited time allowed for its development (three weeks compared to approximately three months). Furthermore, invasion into the lobular compartment with distinguishable ADM (**Figure 5c**), demonstrated the model’s efficacy in recapitulating human PDAC. Next, we quantified immunofluorescence (IF) stains for the basal tumor marker, HMGA2, and used amylase to confirm regions of lobular invasion (**Figure 5d, Supplementary Figure 15**). We selected ROIs based on morphology seen on consecutive H&E stains and quantified HMGA2 in predefined lobular and stromal regions. In line with the autochthonous model and the results from human PDAC, HMGA2 was downregulated in tumor^in_lobules^ compared to tumor^in_stroma^ (**Figure 5e**). Together, these results support a model in which cues from the lobular microniche drive tumor cell phenotypes (**Figure 5f**) and strongly argue against clonal selection as the major source of phenotype heterogeneity.

## Discussion

Reciprocal interactions of tumor cells and their stromal neighbors regulate central functions of the PDAC ecosystem, such as tumor cell differentiation and immune tolerance^16,19^. Manipulating stromal signals can dramatically affect tumor progression, warranting a deep understanding of the cellular interaction partners within the tumor microenvironment^4,5,39^. The current model of PDAC growth is dominated by the dense and heterogeneous stromal reaction surrounding the bulk of the tumor cells^40,41^. In addition to stromal invasion, neurotropism and vascular invasion have been suggested as important anatomic trajectories of PDAC cells^42,43^. Our data confirm that the main interactions of tumor cells are established with the fibrotic stroma, including vessels and nerves. However, in addition to these routes, our results systematically establish another tumor invasion trajectory, the colonization of pancreatic lobules (depicted schematically in **Figure 6**). This is important because, at the leading edge of tumor invasion, PDAC cells meet unique interaction partners, potentially exposing them to a signaling environment distinctly different from that in the stroma, along nerves, or in vessels^21^. Using an automated, unbiased ML approach, we chart these differences on a cellular level and identify cells that have undergone ADM as the main interaction partners of tumor cells that invade lobules.

**Figure 6.**
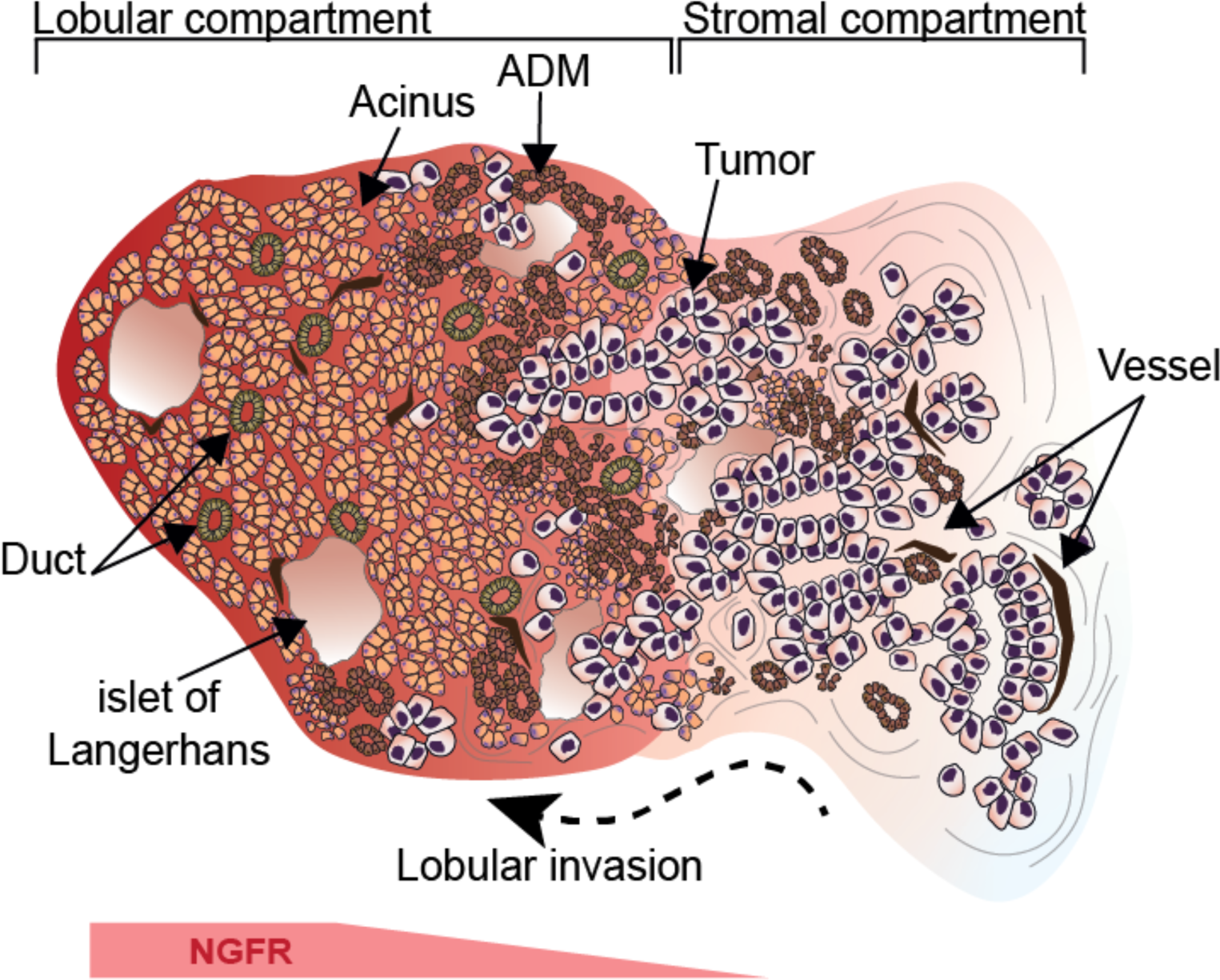
Stromal and tumor cell phenotypes in lobular vs stromal compartments. Illustration of the results. Exocrine cells that undergo acinar-to-ductal metaplasia are the main non-stromal cells adjacent to tumor^in_lobules^, marking the edge of early tumor invasion in the lobules. Nascent stromal cells expressing nerve growth factor receptor (NGFR) in the lobular compartment transition into the NGFR^-^ desmoplastic stroma in the stromal compartment. Gradual loss of NGFR expression develops with increasing lobular atrophy, and is associated recurrently with basal-like tumor cell phenotypes.

ADM is the result of pancreatic injury, for example in pancreatitis^44^. In the context of PDAC, ADM is mainly considered an early event in tumorigenesis, such that reprogramming of acinar cells is thought to be a major initial event in cancer formation^45^. This model is supported by results from genetically engineered mice, which indicate that oncogene activation in acinar cells first leads to ADM, followed by the development of PDAC^12,44,46^. Results of recent single-cell-RNA sequencing studies that have identified ADM cell signatures in PDAC regions have been interpreted according to this model, such that ADM cells present in PDAC are seen as precursor cancer cells, drawing up an intratumoral phylogenetic tree from ADM cells to cancer cells^18^. Our study offers another perspective on these results, as we demonstrate that invading tumor cells in the lobules are invariably associated with ADM in the adjacent exocrine compartment. Therefore, ADM cell gene signatures in advanced PDAC are likely due to adjacency rather than lineage derivation.

We show that PDAC colonization of the lobules is associated with a specific, NGFR^+^/PDGFRA^+^ stroma, while NGFR^+^ stromal cells are rare to absent in desmoplastic regions. NGFR is a marker for benign fibrosis in the pancreas and the liver, where it marks benign-like stroma associated with metastases encapsulation^37^. A previous study has shown in vitro that pancreatic stellate cells temporarily upregulated NGFR expression when co-cultured with tumor cells or tumor cell supernatants^30^. Together with our data, this supports a model where tumor derived factors induce a transient nCAF state in stellate cells, preceding the desmoplastic transformation of the pancreatic parenchyma. While the mechanisms of NGFR need further elaboration, we posit that NGFR is a potential universal marker for very early tumor-induced stromal reprogramming; in our PDAC cases, loss of NGFR in areas of dense stroma was invariably associated with the presence of tumor cells, making NGFR a potentially diagnostic marker to distinguish chronic reactive/inflammatory changes and tumor invasion.

Previously, we have shown that PDAC invading the duodenal wall shifts from a basal to a classical phenotype upon translocation from the submucosa (a stroma-rich compartment) to the mucosa (an epithelial compartment)^34^. Here, we further connect tumor phenotypic heterogeneity with specific tissue compartments inside the pancreas. Cancer cells that gradually colonize the pre-existing pancreatic lobules establish direct cell contact with resident epithelial cells, reminiscent of the mode of PDAC growth in the duodenum^34^ and similar to the replacement-like growth pattern described in the liver^37,47^. Together, the results support a model where tumor-epithelial cell contacts – such as PDAC-ADM, or PDAC-enterocyte^34^ contacts – are associated with a specific tumor cell phenotype. In addition, they might explain the incongruency observed in tumor phenotypes between primary PDAC and its liver metastasis, considering that tumor-epithelial interactions, for example in tumor-hepatocyte contact in liver metastases, might favor the classical phenotype.

Motivated by the stroma-dominant morphology of PDAC, most recent studies on the subtypes and tumor-stromal interactions have focused on studying the tumor cells surrounded by desmoplastic stroma^19,22,40,41^. Many current approaches utilize laser-capture microdissection to enrich tumor cell content prior to RNA sequencing and thus exclude potential areas of lobular invasion. Consequently, tumor cell areas devoid of a desmoplastic stroma have received little attention, and therefore, the characteristics of tumor regions preceding the development of the typical PDAC stroma in human tumors are largely elusive.

Our results indicate local microniche-derived cues rather than clonal selection as major drivers of PDAC subtypes and intratumoral heterogeneity. We propose a model of pancreatic cancer growth in which tumor cell invasion into lobules marks nascent stromal transformation reminiscent of early oncogenic transformation, opening the path for a more nuanced understanding of PDAC evolution in time and space.

## Acknowledgements

The PhD student position of SSö is supported by Karolinska Institutet. This study was supported by The Swedish Research Council (project nr. 2018-02023, to M.G.), The Swedish Society for Medical Research (to M.G.), The Swedish Cancer Society (22 2175 Pj, to M.G.), the Center for Innovative Medicine (to M.G. and J.E.), Cancer Research KI (to M.G. and E.S.), and Radiumhemmet’s research funds (to MG). J.E. was supported by Region Stockholm, the Bengt Ihre Foundation, and Radiumhemmet’s research funds. C.F.M. is supported by The Swedish Society for Medical Research (PD21-0114) and Ruth and Richard Julin’s foundation. Part of this work was carried out at the SciLifeLab CRISPR Functional Genomics unit (CFG) at Karolinska Institutet, funded by Science for Life Laboratory, and Live Cell Imaging core facility/Nikon Center of Excellence at Karolinska Institutet, supported by the KI infrastructure council and Department of Clinical Pathology and Cancer Diagnostics. We gratefully acknowledge Peter Bankhead’s support for staining quantification with QuPath. CFG acknowledges support from the National Genomics Infrastructure, the Swedish National Infrastructure for Computing (SNIC), and the Uppsala Multidisciplinary Center for Advanced Computational Science (UPPMAX). We thank Kjetil Söride for helpful comments on the manuscript and Andrea del Valle for support with animal experiments.

## Material and Methods

### Ethics statement

The work on human samples was approved by the Swedish National Ethical Review Board (*Etikprövningsmyndigheten*, #2020-06115); informed consent was waived. No compensation was provided for participants. The animal experiments were approved by the regional ethics committee the Swedish Board of Agriculture (*Linköpings djurförsöksetiska nämnd*, #00217-2022)

### Patients and slide selection

Patients who underwent surgical resection of a pancreatic ductal adenocarcinoma in 2017 and 2020 were identified in electronic databases at Karolinska University Hospital, Huddinge, Sweden. In clinical routine, one section from each tissue block from the gross sectioning is subjected to H&E staining; based on the tissue content in the H&E sections, one block is further sectioned and stained with an IHC panel, chosen by the pathologist handling the case. Consecutive slides from cases for which 1. IHC stains had been performed, 2. sufficient tumor cellularity for quantification in lobular and stromal compartments, respectively, were selected. Digital WSIs were obtained by scanning with a Hamamatsu NanoZoomer S360 digital slide scanner at x40 magnification.

### Digital semiquantitative scoring of tumor localization

Using both H&E and p53 IHC WSIs of *n* = 5 tumors, the area spanning all tumor cells was annotated with the “Brush” tool in QuPath^26^ version 0.4.3. The tumors were selected to 1. represent different degrees of acinar atrophy in areas of acinar invasion and 2. match the selection of cases included in the machine-learning algorithm workflow described below. All annotations were classified to ‘lobular’, ‘stromal’ or ‘other’. The class ‘other’ was used for tumor located in bile ducts, pancreatic ducts, adipose, nerve, immune clusters or vessels. The areas for each class were then exported for downstream analysis.

### Digital quantification of compartment dependent heterogeneity

Up to *n* = 18 ROIs of the cases included in the quantification cohort were identified blinded to IHC stains (by A.Z.) on stains consecutive to the quantified IHC stains. For example, H&E and IHC stains such as CD146_NGFR, p53-CD34_Caldesmon-CK19, p53-Podoplanin_Caldesmon-Maspin, Chromagranin_CD56, SMAD4 and equilibrative nucleotide transporter (Ent1). The WSI ROIs were dichotomized into tumor^in_lobules^ or tumor^in_stroma^. Tumor^in_lobules^ was defined as tumor cells inside a pancreatic lobule, where lobular cell types or remnants such as endocrine islets of Langerhans were in close proximity. Subsequently, the overlapping regions were identified from the blindly set ROIs in QuPath (S.S.). ROIs were manually annotated with the polygon tool in QuPath, to only include tumor cells in the given compartment, until between 300-1000 cells could be detected in each ROI (**Supplementary Figure 2a**). If needed, the ROIs were expanded until sufficient cell numbers could be included or shifted to the nearest lobular region if the level differences between consecutive sections caused lobular regions in the ROI-selection WSI to be discontinued in the quantification WSI. Next, cell detection and thresholds for the DAB or AP channels for the corresponding stain were applied. The parameters for cell detection and thresholds were adapted to each WSI, determined by visual assessment, listed in **Supplementary Table 1**. The QuPath measurements were exported as .csv files, annotation-wise, including the columns “Image”, “Class”, “Num.Detections”, “Num.Tumor: Negative”, Num.Tumor: Positive”, and “Tumor: Positive %”.

### Generation of supervised learning algorithms for quantification of interactions between tumor cells and non-malignant host cells in lobular and stromal compartments

A nested convolutional neural network-based model was developed with Aiforia Create v 5.5^48^ to identify the different cell and tissue components involved in lobular and stromal invasion on digital WSIs of human PDAC (*n* = 5) stained immunohistochemically with CD146 and NGFR (**Supplementary Figure 10a**). The model was primarily trained to recognize the main structures of the pancreas i.e. acinar cells, ducts, acinar cells undergoing ADM, fibroblasts, vessels, nerves, immune cell clusters, endocrine islets of Langerhans, and tumor cells. First, manual annotations (ground truth) were performed capturing the size and morphological variation of the different structures to increase the robustness of the model (by A.V.). Subsequently, a model was trained, and the output was visually evaluated using Aiforia’s verification and analysis features. Further training annotations were added for structures that resulted in challenges to the model. The process was continued until the model’s performance was deemed adequate upon visual inspection by a specialist pathologist (C.F.M.), and based on verification metrics, with no significant changes in performance observed during the final training rounds. The final model incorporated a total of *n* = 664 training regions and was trained over *n* = 3000 iterations. A detailed description of the training procedure has been given earlier^49^. Details on model hyperparameters and error metrics are given in **Supplementary Table 2.**

### Selection of regions of interest and ML model evaluation

Manually delineated ROIs for different stages of lobular invasion (*n* = 15) and desmoplasia (*n* = 26), were analyzed using the developed ML model. Model accuracy was assessed visually by a pancreas pathologist (C.F.M.). The resulting ML-based annotations were migrated from Aiforia API v 1^50^ into QuPath ^26^, and visually curated one final time, such that major misclassifications were manually curated, for example in areas were tumor and endocrine islets of Langerhans islets or ducts were misclassified (**Supplementary Figure 10a, box 2**). After correction, model accuracy was validated against manual annotations of the tumor cell contacts with the same tissue classes than the model was trained on for lobular invasion (*n* = 3) and tumor in stroma (*n* = 5) in one digital WSI of human PDAC (*n* = 1). In case the contacting tissue with tumor cell was unidentifiable to predetermined classes (i.e. acinar cells, ducts, acinar cells undergoing ADM, fibroblasts, vessels, nerves, immune cell clusters, endocrine islets of Langerhans) due to challenging morphology, caused by, for example, tissue sectioning, the contact was assigned to class ‘other’. After validation, the invasion front for each lobular ROI was annotated, and cell detection was executed within each tumor, ADM, and acinar annotation. Next, the 2D spatial distance between cell detections and ML-based annotations, together with annotations of invasion front, was computed with QuPath. Cell class annotations (e.g. ADM, fibroblast) within 20 µm from the tumor cell detection centroid were considered as contacts since the centroid of tumor cells does not directly interface with the surrounding tissue (**Supplementary Figure 11, box 1**).

### Multiplex RNA ISH

RNA ISH was performed with RNAscope^TM^ HiPlex12 Reagent Kit (488, 550, 650, 750) v2 Standard Assay (ACD, #324400) according to manufacturer’s instructions for FFPE sections (4-5 µm) of patient samples. Pretreatment conditions were adapted to the recommendation for the pancreas and liver. Target retrieval was performed manually as instructed by the manufacturer and FFPE reagent (1:30) was applied. The probes Hs-Krt5-T1 (#547901), Hs-CRP-T2 (#476931), Hs-CPB1-T3 (#569891), Hs-Krt17-T4 (#463661), Hs-SPP1-T5 (#420101), Hs-REG3A-T6 (#312061), Hs-MCAM-T7 (#601731), Hs-Krt19-T8 (#310221), Hs-Anxa10-T9 (#400101), Hs-Spink1-T10 (#469901), Hs-GATA6-T11 (#603131), and Hs-Krt7-T12 (#550151) were used. Images were captured using an inverted confocal Nikon Ti2 microscope in widefield mode with a 20X Plan Apo air objective with an additional 1.5 lens coupled to a Kinetics sCMOS camera. The imaging acquiring details for all RNA targets are summarized in **Supplementary Table 3**.

### Generation of barcoded murine pancreatic adenocarcinoma cells

#### Creating a barcoding library

An existing sgRNA library 51 (cloned by Gibson assembly into pLenti-Puro-AU-flip-3xBsmBI (Addgene, #196709)52 and packaged into lentivirus in HEK-293T (ATCC) using plasmids psPAX2 (a gift from Didier Trono, Addgene, #12260) and pCMV-VSV-G (a gift from Bob Weinberg, Addgene #8454) was used to barcode cells. Briefly, functional titer was estimated from the fraction of surviving KPCT cells after transduction with different amounts of virus and puromycin selection.

#### Creating a barcoded cell population

KPCT cells were transduced with the library virus at an approximate MOI of 0.3 in the presence of 2 µg/ml polybrene. Transduced cells were selected with 2 µg/ml puromycin from day 2 to day 6 post transduction.

#### Genomic DNA, NGS library preparation and NGS

Tumors were homogenized in liquid nitrogen using Freezer/Mill® 6870 (SPEX-sample-prep, 6 cycles, rate 14, 2 min on, 2 min off). The ground tissue was resuspended in Tail buffer (100 mM Tris, 5 mM EDTA, 0.2% SDS, 300 mM NaCl, pH = 8) and genomic DNA was isolated as described previously^53^ with the following modifications: QIAGEN Protease ((#19157, QIAGEN) and PureLink RNAse A ((#12091021, Invitrogen) were used for protein and RNA digest, respectively, and the amount of pre-chilled 7.5 M ammonium acetate (Sigma, # A1542) used for protein precipitation was scaled accordingly. Barcode-containing amplicons were created, sequenced and analyzed as described^52^ using modified primers PCR2_FW acactctttccctacacgacgctcttccgatctcttgtggaaaggacgaaacac PCR3_fw aatgatacggcgaccaccgagatctacac **[i5]** acactctttccctacacgacgctct.

The amplicon was sequenced on Illumina NovaSeq6000, reading 20 cycles Read 1 with custom primer CGATCTCTTGTGGAAAGGACGAAACACCG. NGS data was analyzed with the MaGeCK software, v.0.5.6^54^. Initial barcode complexity was confirmed by including a sample of 40M cells harvested five days after lentiviral barcoding. 77,361 of theoretically 77,441 barcodes were detected. Read counts of all tumour samples were normalized to total read count and rounded to integers. An arbitrary read count threshold of 1,000 was set, and all barcodes with counts below this threshold were removed (in each sample, the removed barcodes contributed less than 0.3% of total reads). Some samples were clonal or near clonal, and in these samples, one or very few barcodes have extremely high read counts, which caused these to bleed through to all other samples due to index hopping, a phenomenon occurring on patterned Illumina flowcells. After filtering for this effect, 931 of 943 barcodes were present in only one of the samples, demonstrating that the samples are independent of each other. Frequencies at which each guide contributes to a sample were calculated based on the normalized total sample read count before filtering.

### Mice

C57BL/6J female mice of 9-11 weeks of age (in house bred but initially obtained from Charles River) were housed in specific-pathogen-free conditions at 12h light/dark cycle. Mice had ad libitum access to standard chow and water, housed at 20-22 °C.

### Ultrasound-guided orthotopic injection

For orthotopic injection, an in-house Pancreatic ductal adenocarcinoma cell line was used (KPCT). Briefly, KPC (*KrasLSL-G12D/+;Trp53LSL-R172H/+;Pdx-Cre*) mice were bred to *B6.Cg-Gt(ROSA)26Sortm9(CAG-tdTomato)Hze/J* mice to generate KPCT mice. Dissociated pieces of KPCT adenocarcinoma from the KPCT mice were provided by Rainer Heuchel. The KPCT cell line was cultured in DMEM/F12 medium (Gibco) with 10% FBS (Sigma-Aldrich) and 1% PenicillinStreptomycin (Sigma-Aldrich) at 37 °C in 5% CO2. At injection d0, the KPCT cells were in passage 18. The orthotopic injection of KPCT cells into the pancreata of mice was performed under anesthesia using ultrasound-guided location of the pancreas with the VEVO 3100 preclinical imaging system (Visualsonics, Toronto, Canada). Mice were anesthetized using isoflurane inhalation. The pain killer, buprenorphine, was administered subcutaneously at 0.05 mg/kg. The fur on the abdomen and left lateral side was gently removed. One mouse at the time was placed in supine position on pre-warmed table in the imaging system, and the pancreas was located with ultrasound. KPCT cells (100 000 cells in max 50 µl sterile PBS) were injected into the pancreas with a 30G needle under ultrasound guidance. Mice were euthanized depending on the welfare assessment either on day 18 after injection day (*n* = 1, M #7), or day 21 (*n* = 3, M #5, #6, M#8). The humane endpoint was defined using a combined score that included the animals’ weight, body posture, physical appearance, behavior, and urine and fecal excretions.

### Tissue processing

Upon euthanasia, the pancreata were harvested, briefly washed in PBS and placed in 4% paraformaldehyde (Sigma Aldrich, #1004965000). After 24h fixation, the tissue was placed in 70% EtOH for at least 24 h followed by paraffin-embedding and sectioning at 4-5 µm thickness.

### Immunofluorescence of murine PDAC sections

The formalin-fixed, paraffin-embedded sections of murine pancreatic ductal adenocarcinoma were baked for 1h at 60°C after which they were deparaffinized, rehydrated, and rinsed in distilled water. Antigen retrieval was performed with pressure kettle in DIVA decloaker (BioCare Medical, #DV2004MX). After blocking with 1% bovine serum albumin, 10% goat serum and 0.1% Triton X-100 for 1 h at room temperature (RT), the sections were incubated sequentially starting with primary antibody at 4°C for overnight, secondary antibody for 1h at RT and lastly, with directly labeled primary antibody for 2 h at RT. All antibodies were diluted to the working concentration with blocking solution. Further details on used antibodies are given in **Supplementary Table 4**. All washing steps were performed with Tris-buffered saline with 0.1% Tween 20. The sections with directly labeled primary antibody were fixed in 4% formaldehyde for 15 min at RT and additional washing step with phosphatase buffer saline before and after fixation and were included. Primary antibodies included HMGA2 (Cell Signaling, #8179, 1:200) and α-amylase (Sigma-Aldrich, #A8273, 1:100). Alexa Fluor 647 goat anti-rabbit IgG (Invitrogen, #A-21245, 1:400) was used as secondary antibody. To directly label the primary antibody, the Zenon Rabbit IgG Labeling Kit was used according to the manufacturer’s instructions (Invitrogen, #Z25302). Counterstaining was done with 4’,6-diamidino-2-phenylindole ([DAPI], Thermo Fisher Scientific, #121101, 1µg/ml) for 15 min at RT. The sections were mounted in ProLong Gold antifade reagent (Invitrogen, #P36924). Images were captured using an inverted confocal Nikon Ti2 microscope in widefield mode with a 20X Plan Apo air objective with an additional 1.5 lens, coupled to a Kinetics sCMOS camera. Imaging details for respective fluorescent probes are summarized in **Supplementary Table 5**.

### m-IF

The multiplex staining was conducted on the Bond RX_m_ autostainer (Leica Biosystems) as described previously^37^. In brief, deparaffinization was achieved with Dewax solution (Leica Bond, #AR9222) followed by an initial antigen retrieval using Epitope retrieval solution 2 (Leica Bond ER2, #AR9640) at 95°C for 30 min. Six target staining cycles were performed in the following antibody order: anti-CD74, anti-PDGFRa, anti-NGFR, anti-p53, anti-ASMA and anti-Vimentin. Details of antibodies and dilutions are provided in **Supplementary Table 6**. Each cycle included a 5 min blocking step with serum free Protein Block (Agilent, #X0909), primary antibody incubation for 30 min, followed by a 10 min incubation with secondary antibody (either ImmPress-mouse HRP, MP-7402, or ImmPress-rabbit HRP, MP-7401, Vector Laboratories) and fluorescent labelling for 10 min using tyramide signal amplification with Opal dyes in the sequence of 690, 620, 520, 570, 480 and TSA-DIG/780 (all Akoya Biosciences), diluted 1:300 in 1X Plus Automation Amplification Diluent (Akoya Biosciences, #FP1609). All steps were carried out at RT. Each staining cycle concluded with an antigen retrieval step with Epitope retrieval solution 1 (Leica Bond ER1, #AR9961) at 95°C for 20 min. Finally, a 5 min DAPI staining (Akoya Biosciences, #FP1490) was performed before mounting with ProLong Diamond Antifade media (Thermo Fisher Scientific, #P36970). Each staining step was followed by subsequent washing steps with Wash solution (Leica Bond, #AR9590).

### Staining quantification of HMGA2+ murine tumor cells

On sections of murine PDAC (*n* = 4) with HMGA2 and amylase IF costain, ROIs of stromal invasion (*n* = 16) and lobular growth (*n* = 24) were delineated. The definition of the ROIs was done by one researcher (M.G.) on H&E-stained sections, blinded to IF. Another researcher (A.V.) performed the quantification of the matched IF images. The HMGA2 channel was kept off while delineating the corresponding IF regions. For each ROI, cell detection based on nuclear DAPI staining and positive cell detection based on HMGA2 mean nuclear intensity in the AP channel were called. Nuclear mean intensities for HMGA2 ranged between 121-3860 arbitrary units (AU). Intensity values over 300 AUs were considered as positive. An example of positive cell detection is given in **Supplementary Figure 15**. Quantitation data were exported in tabular format for downstream analysis.

### IHC-based quantification of stromal stains

For the same human PDAC sections double stained with IHC for CD146 (DAB-brown chromogen) and NGFR (AP-red chromogen), excluding case 7 (*n* = 4) due to end-stage morphology and negativity for NGFR stain, NGFR intensity of ROIs for lobular invasion (*n* = 12 ROIs) and desmoplasia (*n* = 20 ROIs) were analyzed. For each cell detection inside ML-derived acinar, ADM and tumor annotations, the mean intensity of the AP channel based on 0.1 µm pixel size inside the surrounding circular area (diameter 50 µm) was measured with the “Add intensity features” -command with QuPath (**Supplementary Figure 11 boxes 2&3**). Only cell detections directly interacting with ML-derived fibroblasts annotations (i.e. distance from cell detection centroid to the closest annotation was less than 20 µm) were included for downstream analysis. All recorded negative intensity values were set to zero.

### m-IF based quantification of stromal stains

The circumference of tumor clusters was annotated with the polygon tool in QuPath, using p53+ nuclei for tumor identification. For each case, n = 6 annotations for tumorin_lobules and n = 7-10 annotations for tumorin_stroma were drawn. Three layers of the area in immediate proximity to the tumor annotation were expanded with 15 µm increments with the “Expand annotations” tool in QuPath (**Supplementary Figure 12a**). To analyze the stroma cells present in the different compartments, larger lobular areas (n = 53) or stromal areas (n = 40) were annotated (**Supplementary Figure 13a-b**) Cell detection was performed within the annotations with a subsequent removal of cells with high circularity to enrich for stroma cell content. A cell classifier was trained to recognize cells that were positive or negative by visual assessment and applied to all cell detections. In the subsequent data analysis, any non - stromal residual cells were removed by excluding cells detected as p53+ by the classifier.

### Statistics and reproducibility

All downstream analyses were conducted using RStudio releases ‘Bird Hippie’ or ‘Desert Sunflower’ and R statistical program v 4.2.1 and 4.2.2^55^. Dplyr v 1.1.3^56^, tidyr v 1.3.0^57^ tidyverse v 2.0.0^58^, stats v 4.1.2, were used for data wrangling. Summary statistics were performed with BioGenerics v 0.40.0^59^. Data from the compartment-dependent heterogeneity and multiplex-immunofluorescence pipeline were analyzed with Benjamini-Hochberg-corrected, two-sided Wilcoxon rank sum test or Kruskal-Wallis test with Dunn’s test as a post-hoc from rstatix v 0.7.2. Wilcoxon rank sum test of HMGA2 positive murine pancreatic ductal adenocarcinoma cells between acinar and septal invasion was performed with rstatix v 0.7.2^60^. Visualizations were done with ggalluvial v 0.12.5^61^, ggplot2 v 3.4.3^62^, ggpubr v 0.6.0^63^, ggforce v 0.4.1^64^, ggbreak v 0.1.2^65^, circlize v 0.4.15^66^, ComplexHeatmap v 2.10.0^67^, and enhanced using color scales from RColorBrewer v 1.1.3^68^ and wesanderson v 0.3.7^69^. No data imputation was performed. No statistical method was used to predetermine sample size. Where data are presented as Box-and-Whisker plots, the mean (diamond), the median (line), the interquartile range (IQR, box), minimum and maximum values within 1.5 times IQR from the first and third quartile (whiskers) are shown. In **Supplementary Table 7**, the corresponding tumor ID labels shown in plots to representative images are stated.

---

**Supplementary Figure 1.**
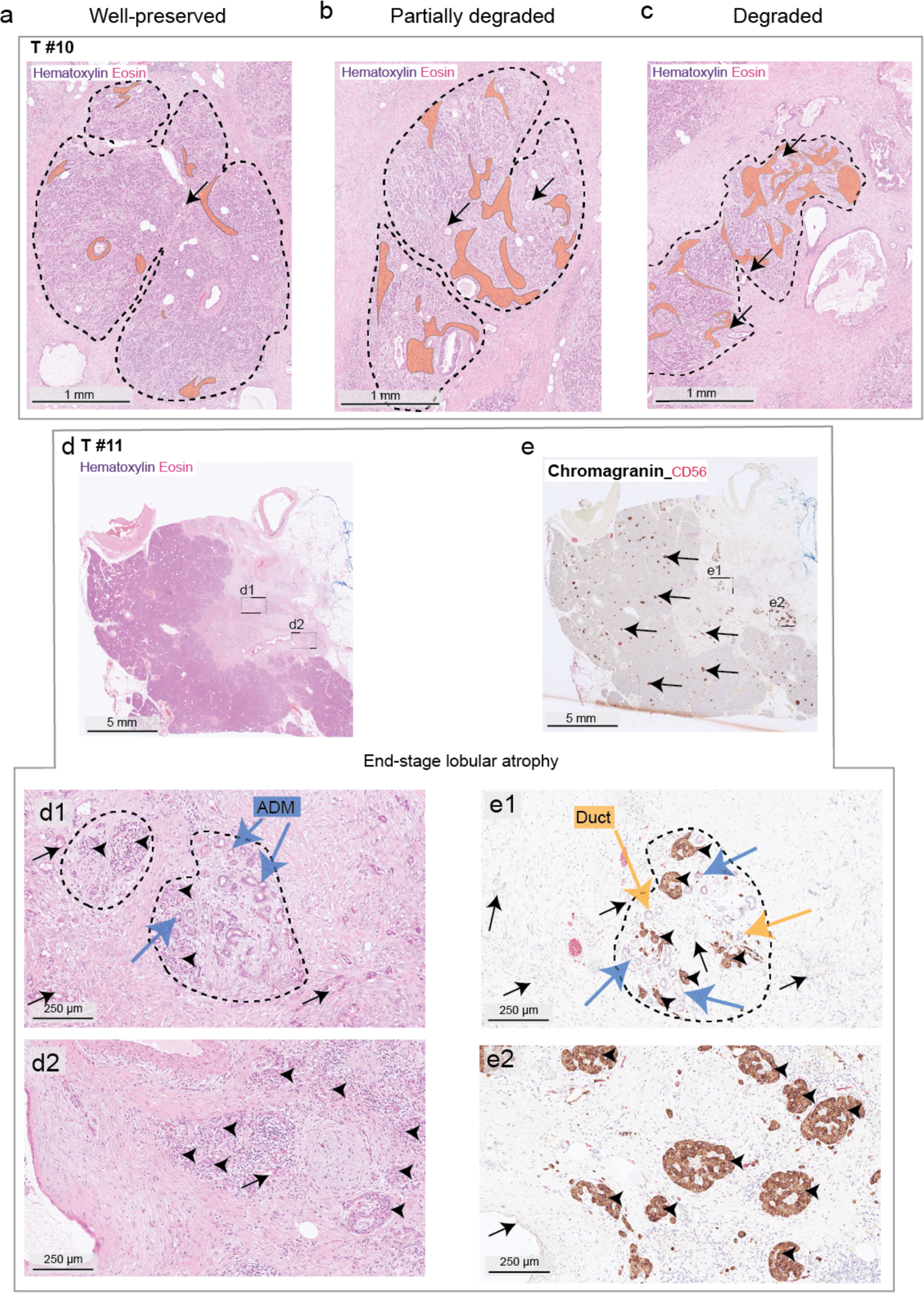
Decay of the pancreatic parenchyma during tumor expansion. **a)-c)** Representative images of increasing lobular degradation, accompanied with stromal expansion (light orange overlay) and tumor invasion, hematoxylin & eosin (H&E) stains: Well-preserved lobule (**a**); as a lobule is degraded (**b**), the parenchyma is dissociated with more pronounced stroma, eventually resulting in complete stroma-transformation (**c**). **d)** H&E and chromogranin A stain (brown, endocrine cells) on partial (**d1** and **d2**) or end-stage (**d2** and **e2**) lobular degradation. Blue arrows: Acinar-to-ductal metaplasia, yellow arrows: ductular remnants, small black arrows: endocrine islets of Langerhans, black arrows with tail: tumor. Dashed line: pancreatic lobules. Representative of *n* = 31 tumors.

**Supplementary Figure 2.**
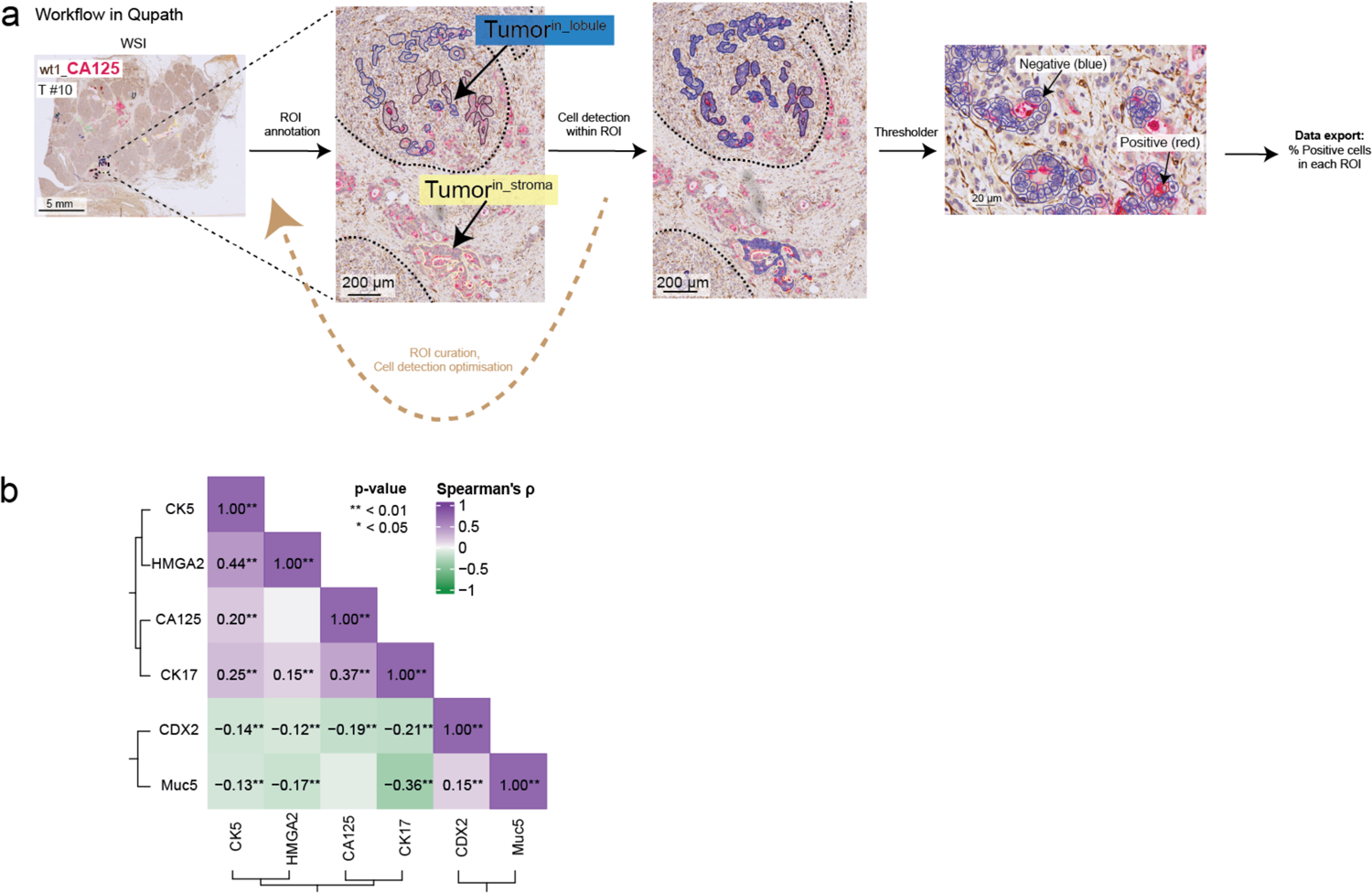
QuPath annotation workflow in compartment-dependent heterogeneity. **a)** Overview of the annotation workflow in QuPath, here exemplified with a whole-slide image (WSI) stained for Carbohydrate antigen (CA)125 (red, cytoplasmic) costained with a stromal marker, WT1 transcription factor ([WT1] brown, cytoplasmic). Up to *n* = 18 ROIs were annotated for each WSI, which were dichotomized to represent tumor in the lobular or stromal compartment. **b)** Correlogram of all markers quantified with immunohistochemistry. Significance levels indicated are * = 0.05, ** = 0.01.

**Supplementary Figure 3.**
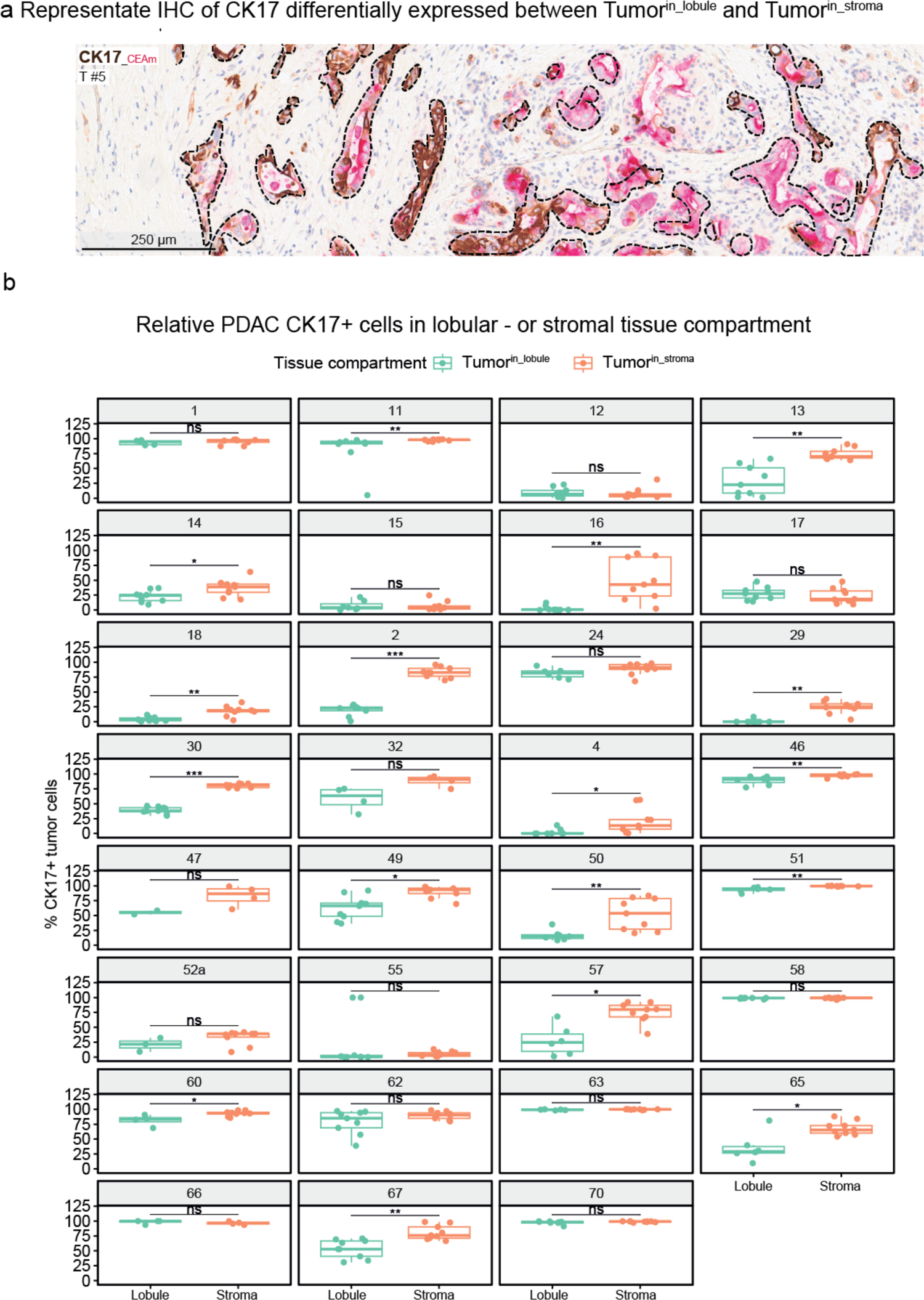
CK17 is more frequently expressed by tumors in stroma than in pancreatic lobules. **a)** Representative immunohistochemistry of the basal-like marker Cytokeratin (CK)17 (brown, cytoplasmic, costained with monoclonal Carcinoembryonic antigen ([CEAm] red, cytoplasmic) differentially expressed between lobular– and stromal invasion. Stroma toward left, pancreatic lobule toward right. **b)** Unpaired Wilcoxon rank sum test with Benjamini-Hochberg corrected for multiple testing of quantifications of each case clinically stained with CK17_CEAm. Up to nine regions of interest (ROIs) per compartment (that is, up to *n* = 18 ROIs in total per case) were manually annotated in QuPath to include tumor cells in the respective compartment, and subsequently thresholded for the brown DAB channel to stratify CK17 positive and negative tumor cells. Each dot represents the fraction of CK17+ tumor cells in one individual ROI. Representative image of and data from *n* = 31 tumors. ns: non-significant, * = 0.05, ** = 0.01, *** = 0.001, **** = 0.0001.

**Supplementary Figure 4.**
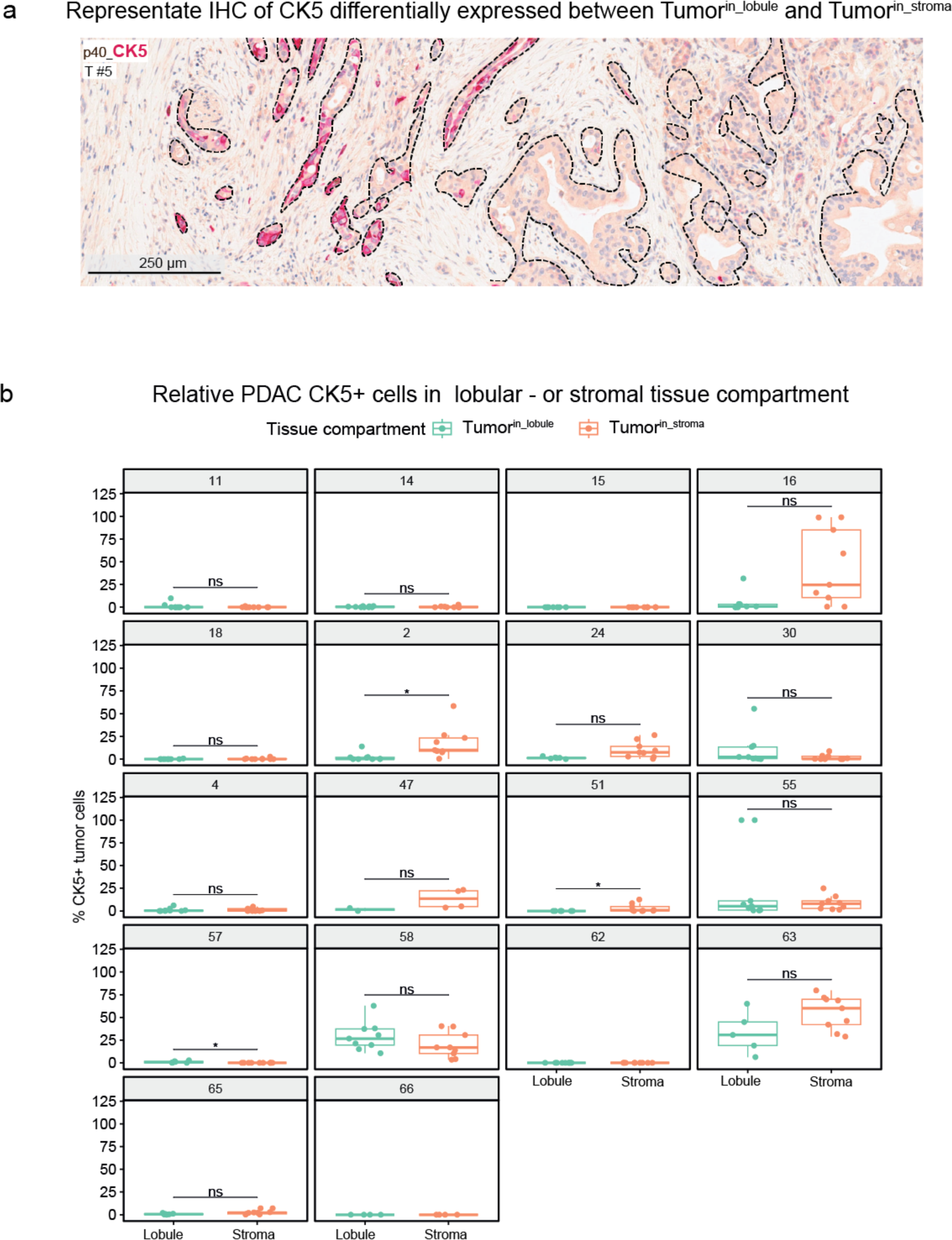
CK5 is more frequently expressed by tumors in stroma than in pancreatic lobules. **a)** Representative immunohistochemistry of the basal-like marker Cytokeratin (CK)5 (red, cytoplasmic) costained with p40 (brown, nuclear) differentially expressed between lobular – and stromal invasion. Stroma toward left and pancreatic lobule toward right. **b)** unpaired Wilcoxon rank sum test, with Benjamini-Hochberg correction for multiple testing of quantifications of each case clinically stained with p40_CK5. Up to nine regions of interest (ROIs) per compartment (that is, up to *n* = 18 ROIs in total per case) were manually annotated in QuPath to include tumor cells in the respective compartment, and subsequently thresholded for the red AP channel to stratify CK5 positive and negative tumor cells. Each dot represents the fraction of CK5+ tumor cells in one individual ROI. Representative image of and data from *n* = 31 tumors. ns: non-significant, * = 0.05, ** = 0.01, *** = 0.001, **** = 0.0001.

**Supplementary Figure 5.**
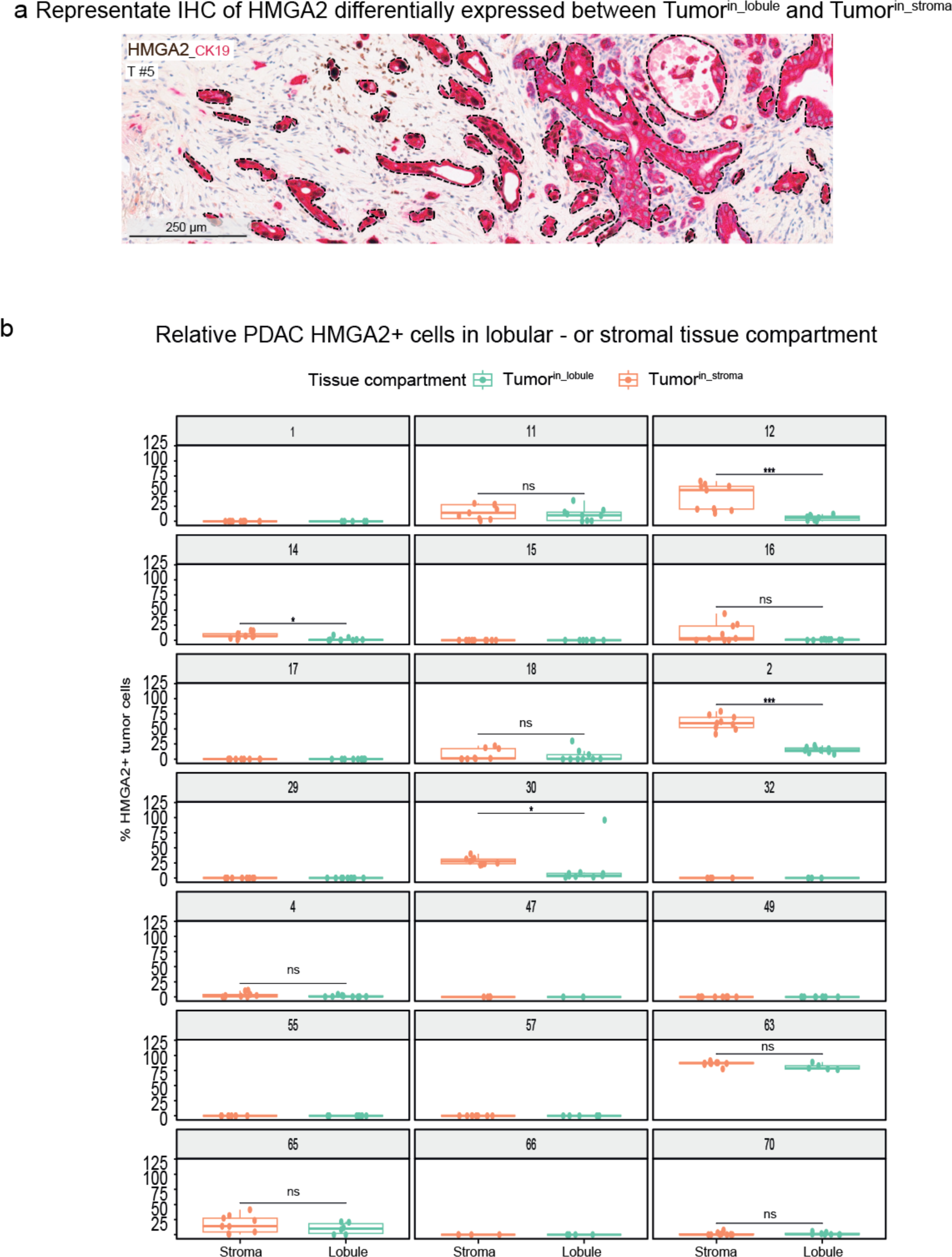
HMGA2 is more frequently expressed by tumors in stroma than in pancreatic lobules. **a)** Representative immunohistochemistry of the basal-like marker high mobility group AT-hook 2 (HMGA2, brown, nuclear costained with Cytokeratin (CK)19, red) differentially expressed between lobular– and stromal invasion. Stroma toward left and pancreatic lobule toward right. **b)** unpaired Wilcoxon rank sum test, with Benjamini-Hochberg correction for multiple testing of quantifications of each case clinically stained with HMGA2_CK19. Up to nine regions of interest (ROIs) per compartment (that is, up to *n* = 18 ROIs in total per case) were manually annotated in QuPath to include tumor cells in the respective compartment, and subsequently thresholded for the brown DAB channel to stratify HMGA2 positive and negative tumor cells. Each dot represents the fraction of HMGA2+ tumor cells in one individual ROI. Representative image of and data from *n* = 31 tumors. ns: non-significant, * = 0.05, ** = 0.01, *** = 0.001, **** = 0.0001.

**Supplementary Figure 6.**
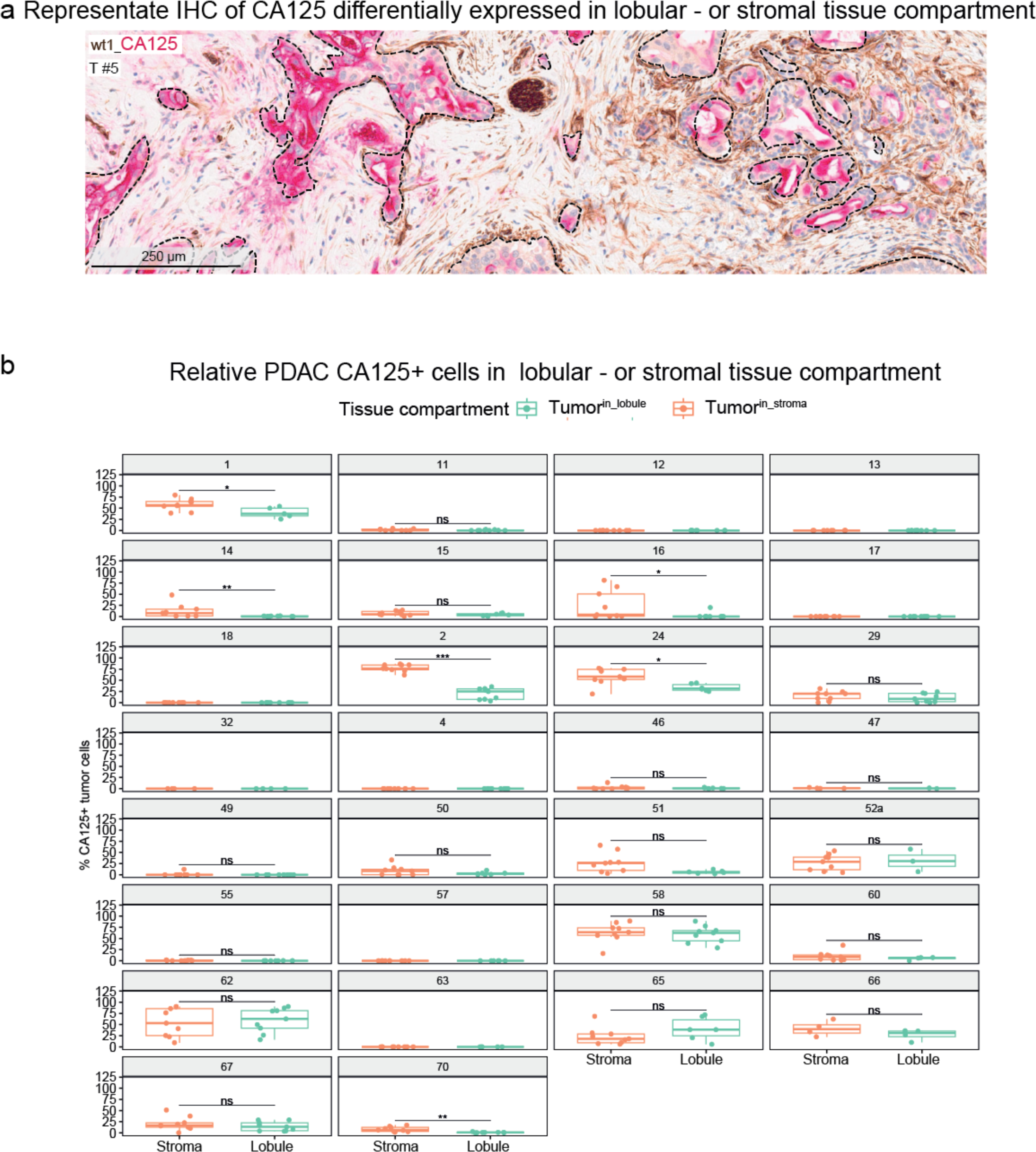
CA125 is more frequently expressed by tumors in stroma than in pancreatic lobules. **a)** Representative immunohistochemistry of the basal marker Carbohydrate antigen (CA)125 (red, cytoplasmic, costained with wt1, brown) differentially expressed between lobular– and stromal invasion. Stroma presented toward left and pancreatic lobule toward right. **b)** unpaired Wilcoxon rank sum test, with Benjamini-Hochberg correction for multiple testing of quantifications of each case clinically stained with wt1_CA125. Up to nine regions of interest (ROIs) per compartment (that is, up to *n* = 18 ROIs in total per case) were manually annotated in QuPath to include tumor cells in the respective compartment, and subsequently thresholded for the red AP channel to stratify CA125 positive and negative tumor cells. Each dot represents the fraction of CA125 + tumor cells in one individual ROI. Representative image of and data from *n* = 31 tumors. ns: non-significant, * = 0.05, ** = 0.01, *** = 0.001, **** = 0.0001.

**Supplementary Figure 7.**
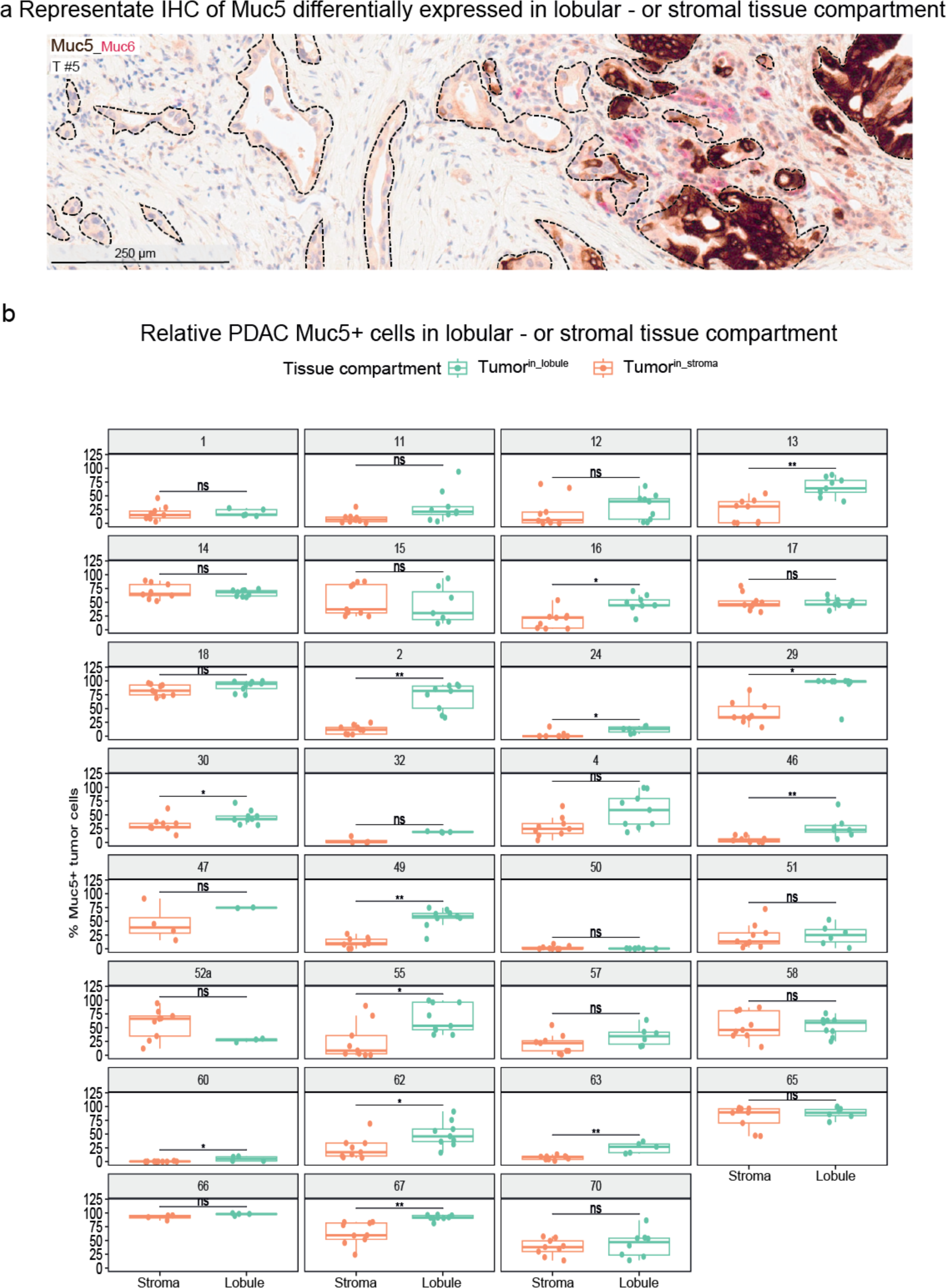
MUC5AC is more frequently expressed by tumors in pancreatic lobules than in stroma. **a)** Representative immunohistochemistry of the classical marker mucin (MUC) 5 subtypes A and C ([MUC5AC], brown, cytoplasmic), costained with MUC6 (red, cytoplasmic) differentially expressed between lobular – and stromal invasion. Stroma toward left and pancreatic lobule toward right. **b)** unpaired Wilcoxon rank sum test, with Benjamini-Hochberg correction for multiple testing of quantifications of each case clinically stained with MUC5AC_MUC6. Up to nine regions of interest (ROIs) per compartment, (that is, up to *n* = 18 ROIs in total per case) were manually annotated in QuPath to include tumor cells in the respective compartment, and subsequently thresholded for the brown DAB channel to stratify MUC5AC positive and negative tumor cells. Each dot represents the fraction of MUC5AC+ tumor cells in one individual ROI. Representative image of and data from *n* = 31 tumors. ns: non-significant, * = 0.05, ** = 0.01, *** = 0.001, **** = 0.0001.

**Supplementary Figure 8.**
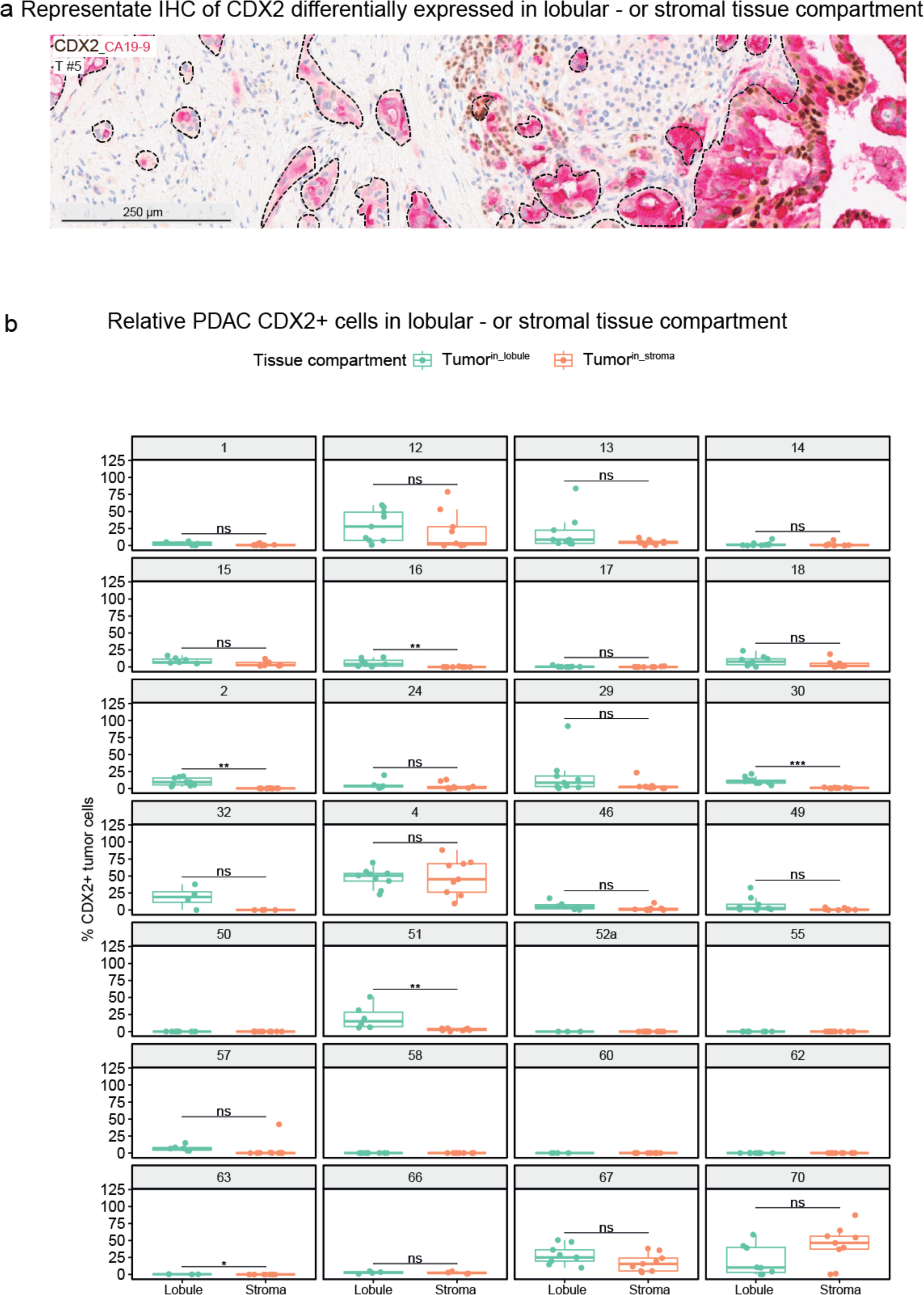
CDX2 is more frequently expressed by tumors in pancreatic lobules than in stroma. **a)** Representative immunohistochemistry of the classical marker caudal type homeobox 2 (CDX2) (brown, nuclear, costained with Carbohydrate antigen (CA)19-9, red) differentially expressed between lobular– and stromal invasion. Stroma presented toward left and pancreatic lobule toward right. **b)** unpaired Wilcoxon rank sum test, BH-corrected for multiple testing of quantifications of each case clinically stained with CDX2_CA19-9. **b)** unpaired Wilcoxon rank sum test, with Benjamini-Hochberg correction for multiple testing of quantifications of each case clinically stained with CDX2_CA19-9. Up to nine regions of interest (ROIs) per compartment, (that is, up to *n* = 18 ROIs in total per case) were manually annotated in QuPath to include tumor cells in the respective compartment, and subsequently thresholded for the brown DAB channel to stratify CDX2 positive and negative tumor cells. Each dot represents the fraction of CDX2+ tumor cells in one individual ROI. Representative image of and data from *n* = 31 tumors. ns: non-significant, * = 0.05, ** = 0.01, *** = 0.001, **** = 0.0001.

**Supplementary Figure 9.**
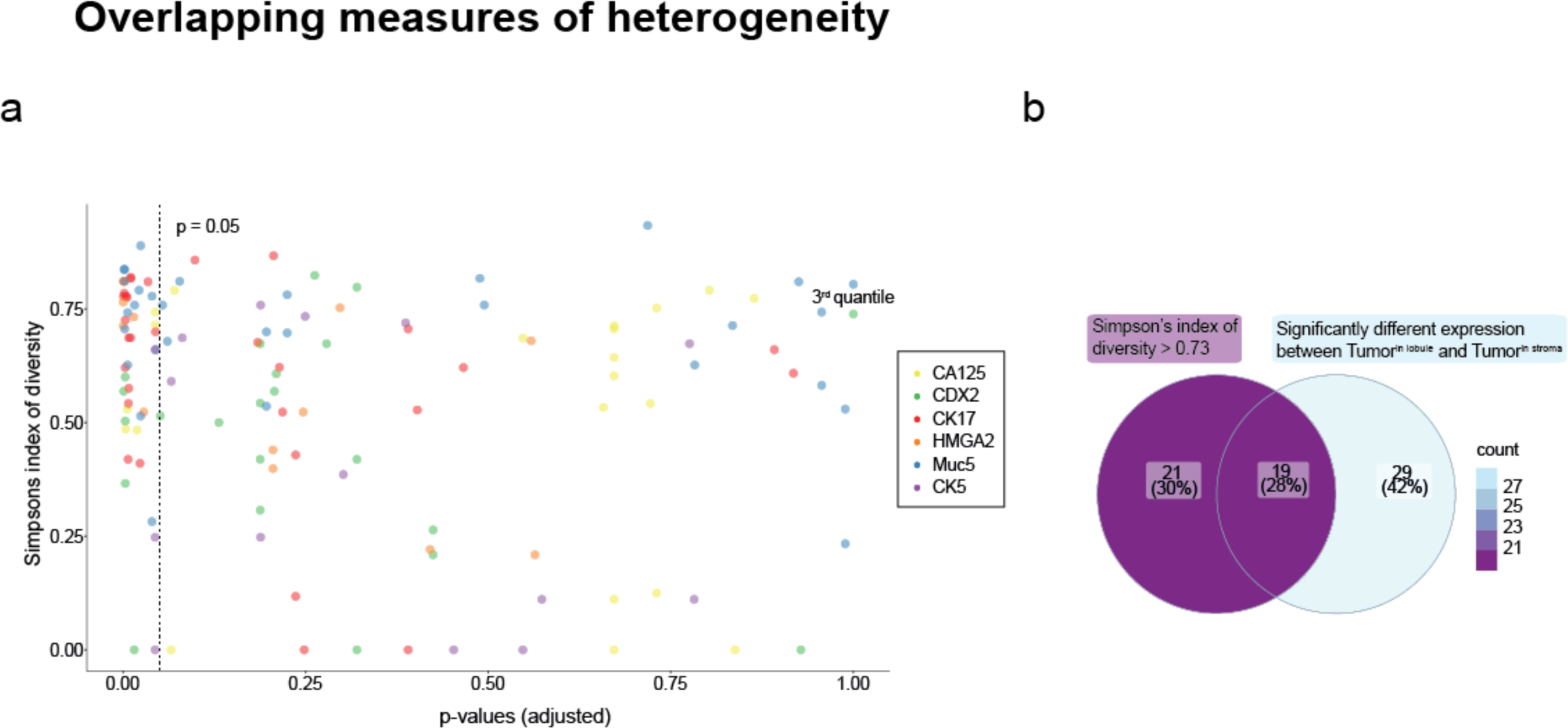
Spatially dependent heterogeneity explains some but not all intratumoral heterogeneity in PDAC. The Simpson’s index of diversity here represents the probability to retrieve two random regions of interest from each stain and tumor that are in different intensity categories, irrespective of which tissue compartment they are localized within. **a)** p-values from Wilcoxon test **(Supplementary Figure 3-8**) against the Simpson’s index of diversity. Dots within the upper left dotted square represent cases and stains significantly expressed between lobular and stromal compartment and with high overall heterogeneity. **b)** Venn diagram of the same data as (**a)** showing cases and stains that have both high spatially dependent and overall heterogeneity account for 28% of all high-heterogeneity instances. Data from *n* = 31 tumors.

**Supplementary Figure 10.**
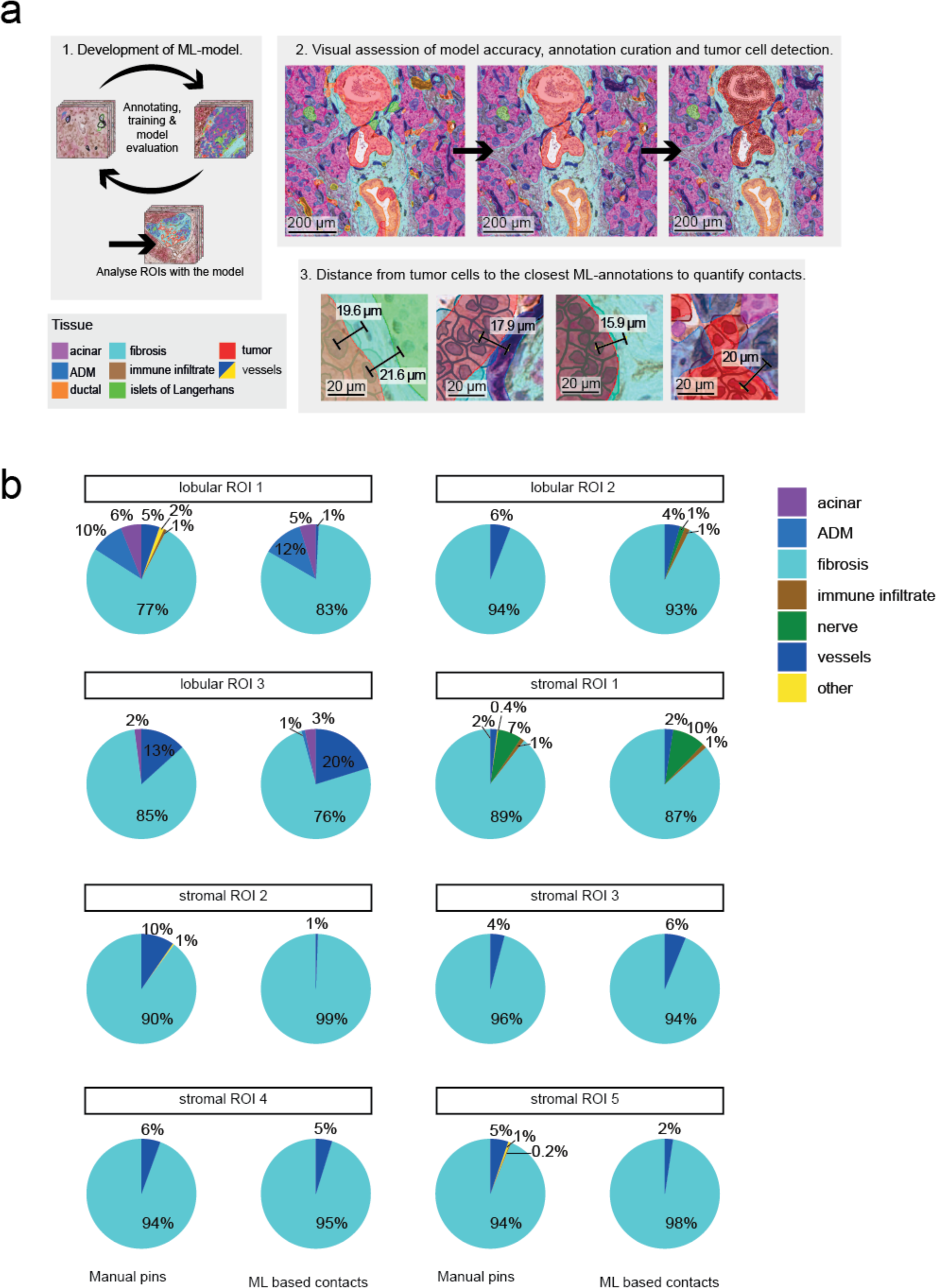
Workflow of machine learning model construction for interaction mapping and validation to manual annotations. **a)** Schematic overview of model construction: (1) curation and analysis pipeline (2) quantitation of tumor contacts to all other determined tissue types based on distance. Distances < 20 µm from tumor cell centroids to each closest tissue class were considered as contact, as exemplified in (3). **b)** Fractions of contacts detected between tumor and each tissue class, either by manually annotating each contact (left) or determined with machine-learning based annotations (right). Data from *n* = 1 tumor. The class ‘other’ comprises manually pinned contacts identifiable as a cell, characterized by the presence of visible nucleus but due to challenging morphology, caused, for example, by tissue sectioning, could not be assigned a class.

**Supplementary Figure 11.**
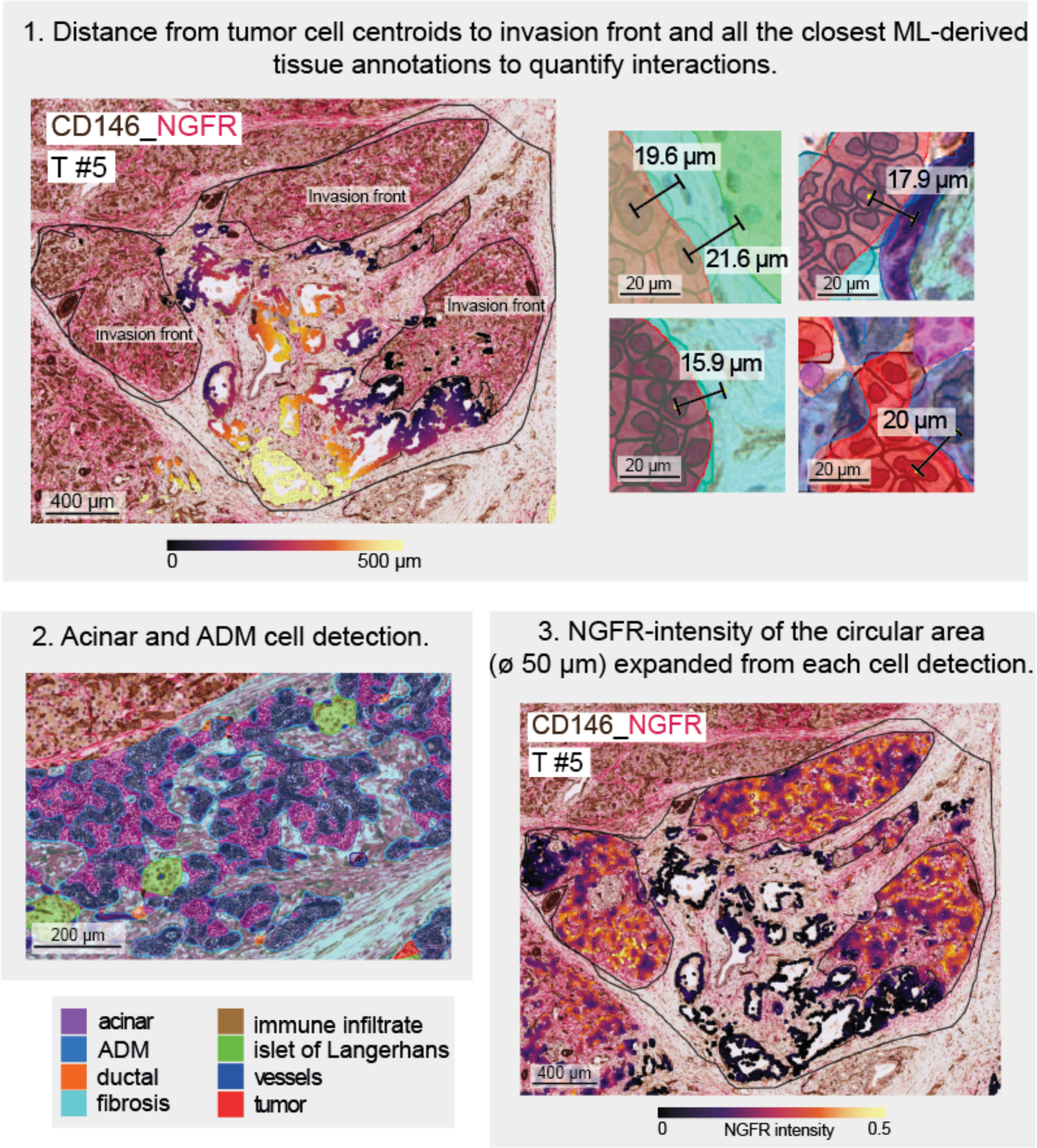
Workflow for spatial interaction mapping and quantification of nerve growth factor receptor (NGFR) expression in the lobules. On duplex immunohistochemistry of cluster of differentiation (CD)146 and NGFR, the distances from the invasion front to tumor cell detections were measured, and contacts with tumor cells were quantified with machine learning (ML) based annotations. Distance < 20 µm was considered as contact, as exemplified (1). For quantification of NGFR gradient, cells were detected inside ML annotated ‘acinar’ and acinar cells undergoing acinar-to-ductal-metaplasia (‘ADM’) classes (2). NGFR stain intensity of expanded area (ø50 µm) from each cell detection (3).

**Supplementary Figure 12.**
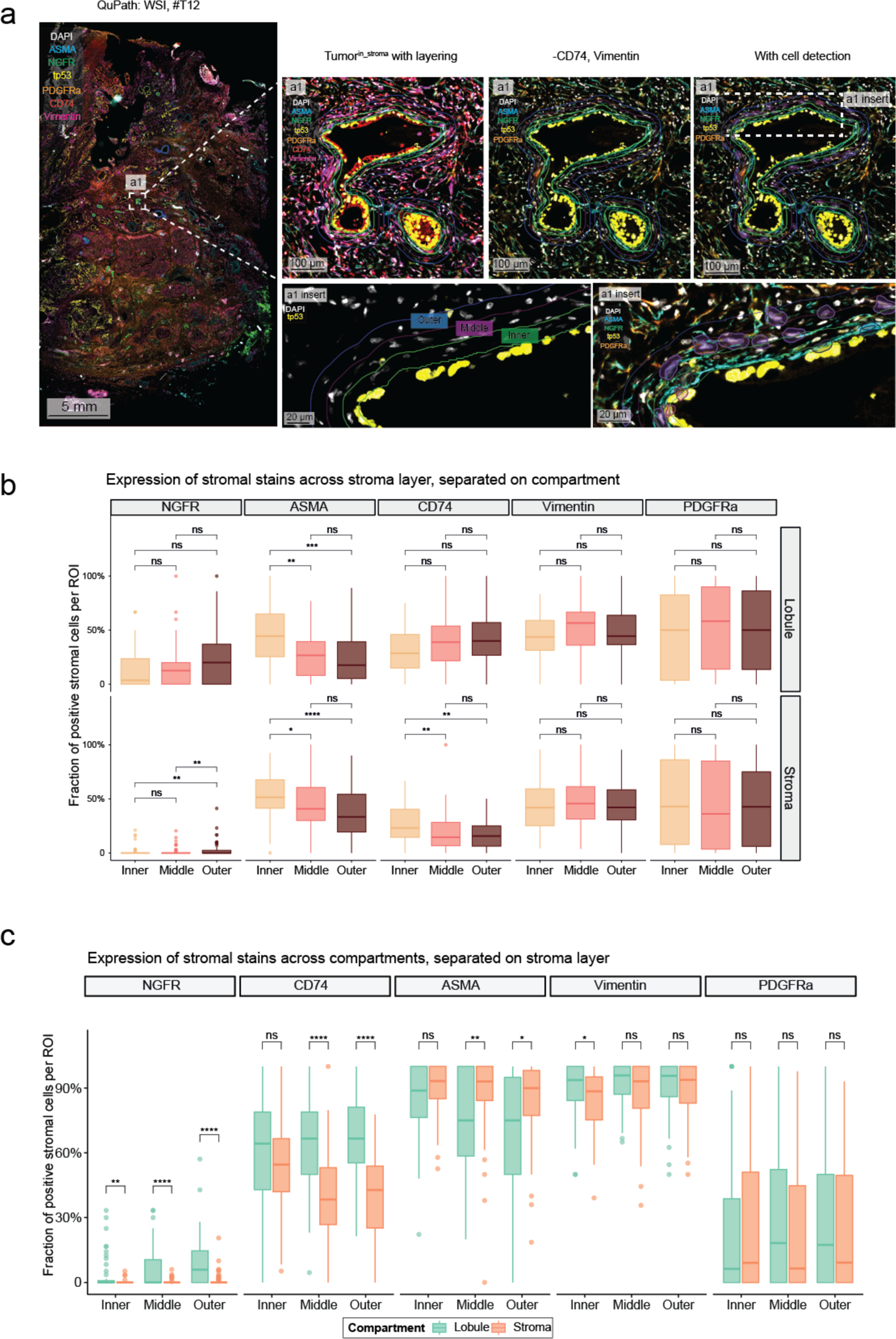
Differential expression of stromal cells with increasing distance from tumor cells. **a)** Overview of the stroma layering approach in QuPath. p53+ tumor nests (yellow, nuclear) were annotated, from where three stromal layers were created.; inner; closest tumor, middle, and outer; furthest away from tumor. **b)** Differential distribution of stroma cells expressing nerve growth factor receptor (NGFR), alpha-smooth muscle actin (ASMA), cluster of differentiation (CD)74, Vimentin, and platelet-derived growth factor receptor alpha (PDGFRa) at three distance layers from tumor nests, separated on tumor^in_lobules^ and tumor^in_stroma^. Kruskal-Wallis rank sum test, with post-hoc Dunn’s test for pairwise multiple comparisons. Related to Figure 4e. **c)** Dichotomized stratification of stroma cells in present next to tumor in lobules and stromal regions at various distances to tumor cell nests. Unpaired two-tailed Wilcoxon rank sum test, with Benjamini-Hochberg correction for multiple testing. Related to Figure 4f. **b)**, **c)**: Data from *n* = 8 tumors, with *n* = 222 stromal regions of interest ([ROIs], *n* = 74 per stroma layer), and *n* = 141 lobular ROIs (*n* = 47 per stroma layer).

**Supplementary Figure 13.**
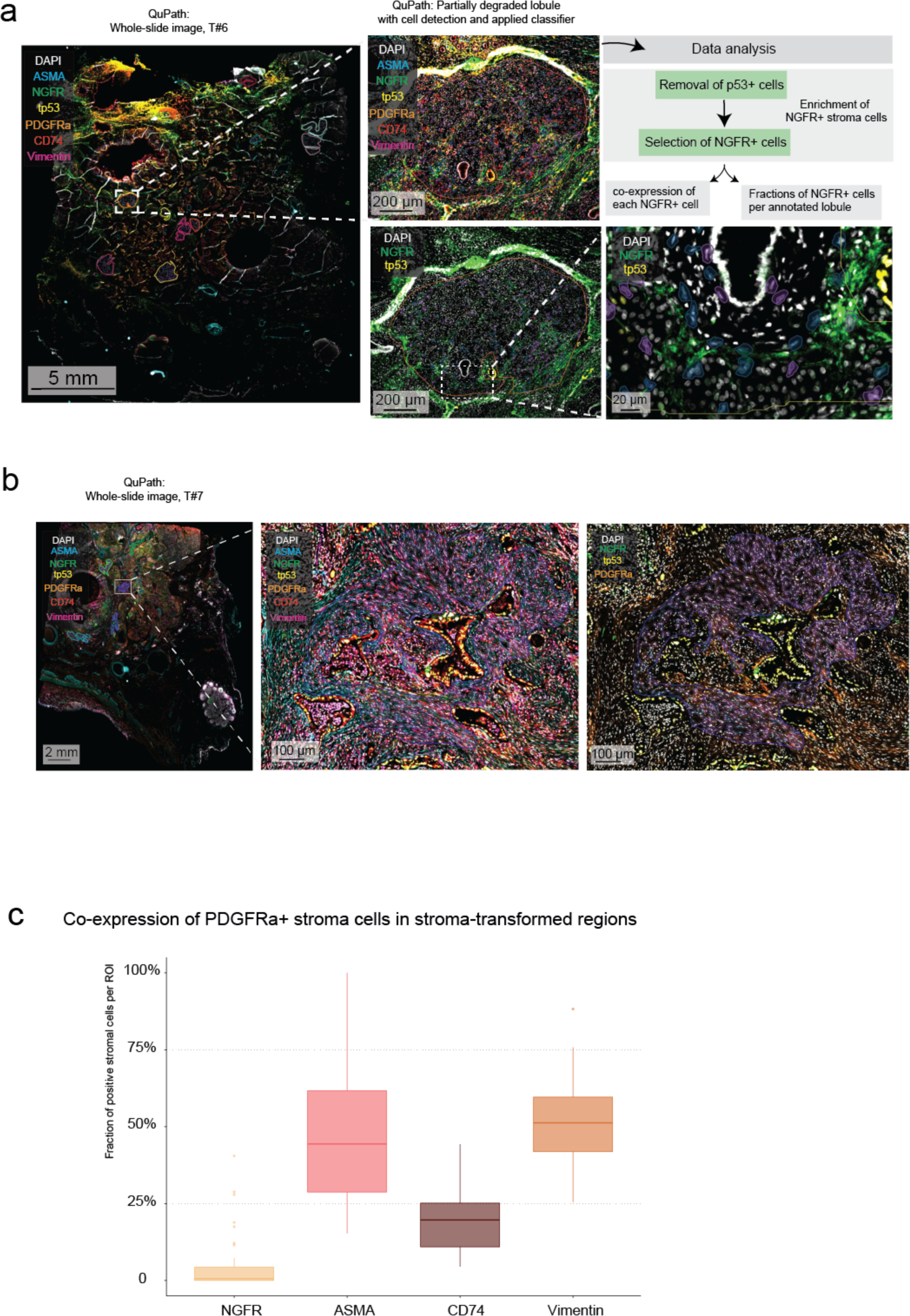
Stromal cells present in the lobular and stroma-transformed compartments have separate expression profiles. **a)** Representative image and explanation of the workflow to analyze nerve growth factor receptor (NGFR)+ stroma cells present in pancreatic lobules. Whole-lobule annotations were manually drawn, in which cells were detected. Cells with high nuclear circularity and p53 positivity were excluded. All cells were subject to a classifier, denoting the expression status of each stain. **b)** Representative image and explanation of the workflow to analyze the platelet-derived growth factor receptor alpha (PDGFRa+) stroma cells present in desmoplastic, stroma-transformed regions. Stromal annotations were manually drawn at sizes comparable to the lobular annotations in (**a)**, in which cells were detected. Cells with high nuclear circularity and p53 positivity were excluded. All cells were subject to a classifier, denoting the expression status of each stain. **c)** The co-expression of PDGFRa+ stroma cells in desmoplastic, stroma-transformed regions). Data from *n* = 8 individual tumors, with *n* = 40 regions of interest in total.

**Supplementary Figure 14.**
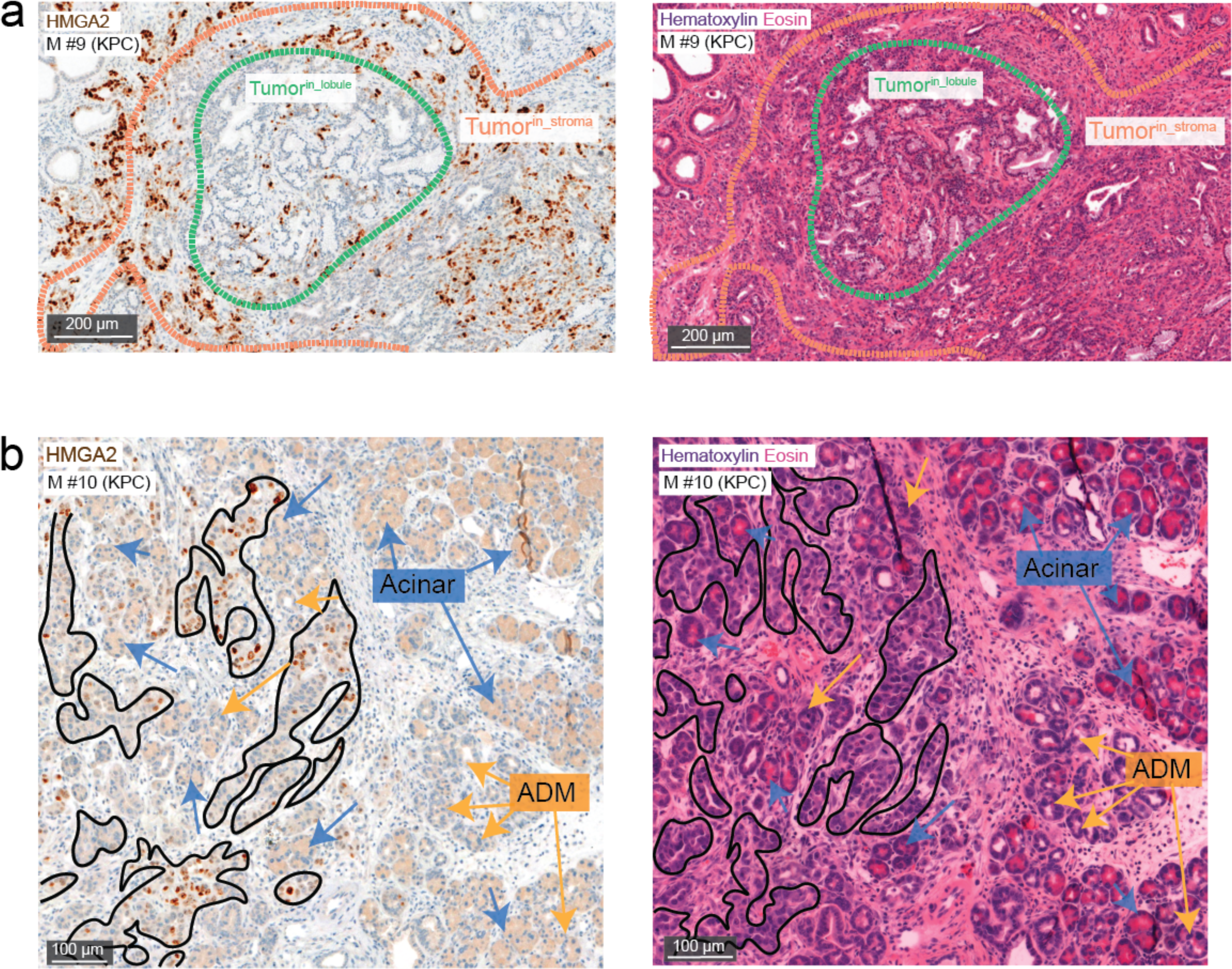
Representative images of immunohistochemistry staining of HMGA2 and H&E from KPC mice. **a)** Expression of high mobility group AT-hook 2 (HMGA2, left panel) by tumor cells in the stromal compartment (orange dotted line) and lobular compartment (green dotted line), and a consecutive hematoxylin & eosin H&E (right panel) **b)** In higher magnification of HMGA2 stain (left panel) and a consecutive H&E (right panel), the tumor cells in lobular compartment (black filled line) advance next to the exocrine cells (acinar, blue arrows; and acinar-to ductal metaplasia (ADM), yellow arrows). a), b) Representative of *n* = 4 tumors.

**Supplementary Figure 15.**
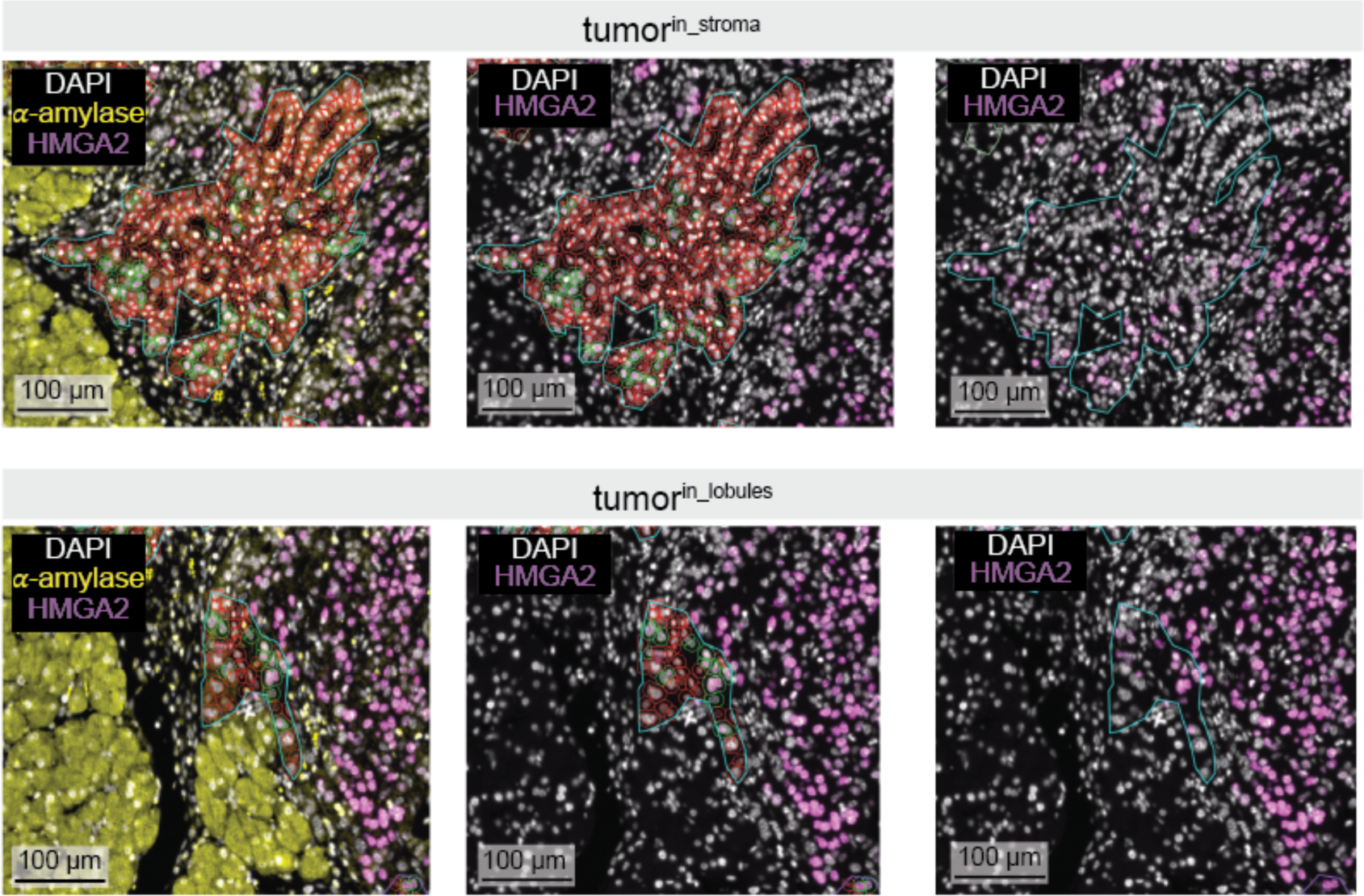
Examples of HMGA2^+^ and HMGA2^-^ tumor cells detected and classified in QuPath. Representative immunofluorescence of high mobility group A2 (HMGA2; pink, nuclear) and α-amylase (yellow, cytoplasmic) of a mouse tumor from the orthotopic injection model of stromal invasion (tumor^in_stroma^, upper panel) and lobular invasion (lower panel). Lines delineate quantified regions of interest, red detection objects: HMGA2-tumor cells, green detection objects: HMGA2+ tumor cells. Representative of *n* = 4 tumors.

**Table S1:**
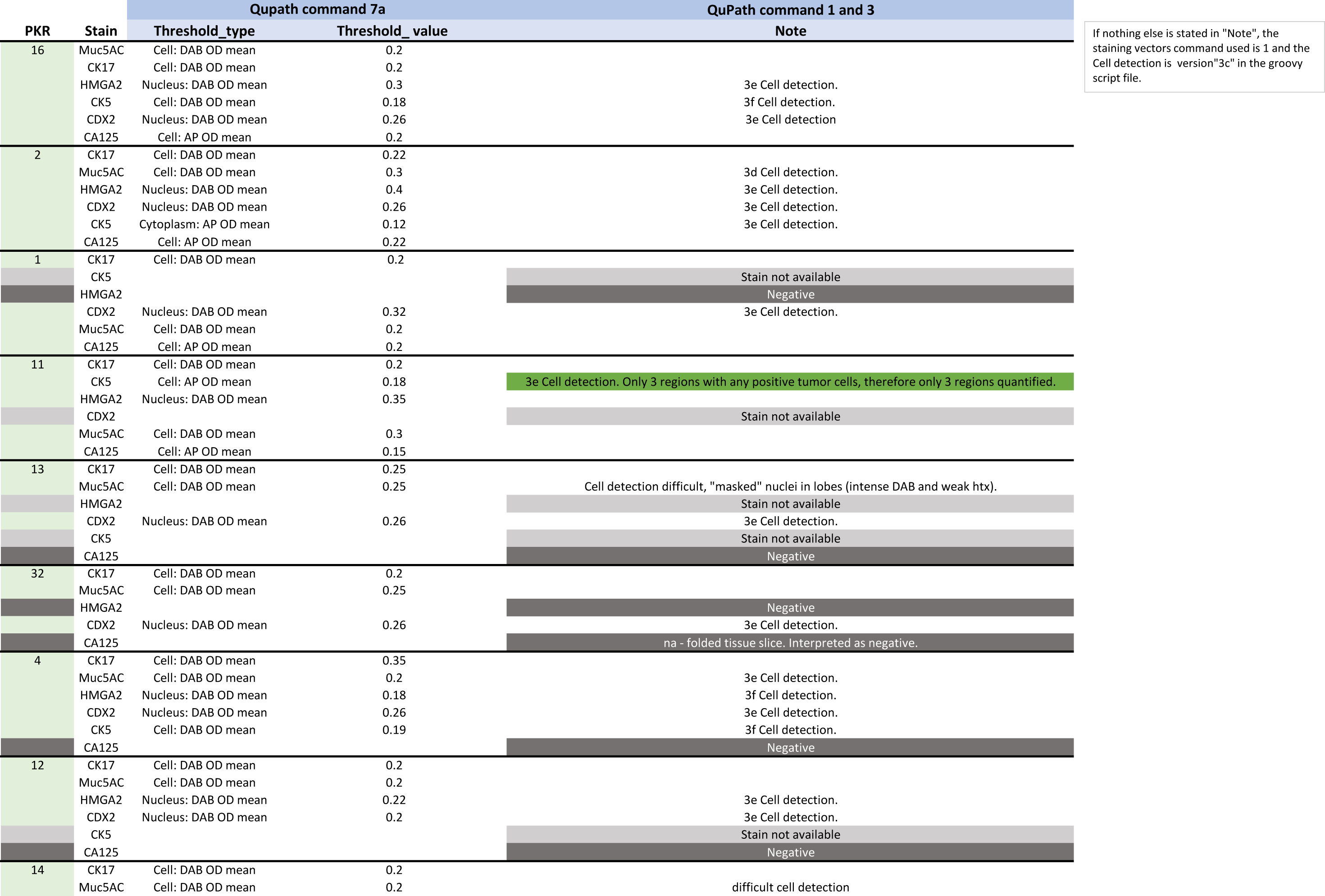
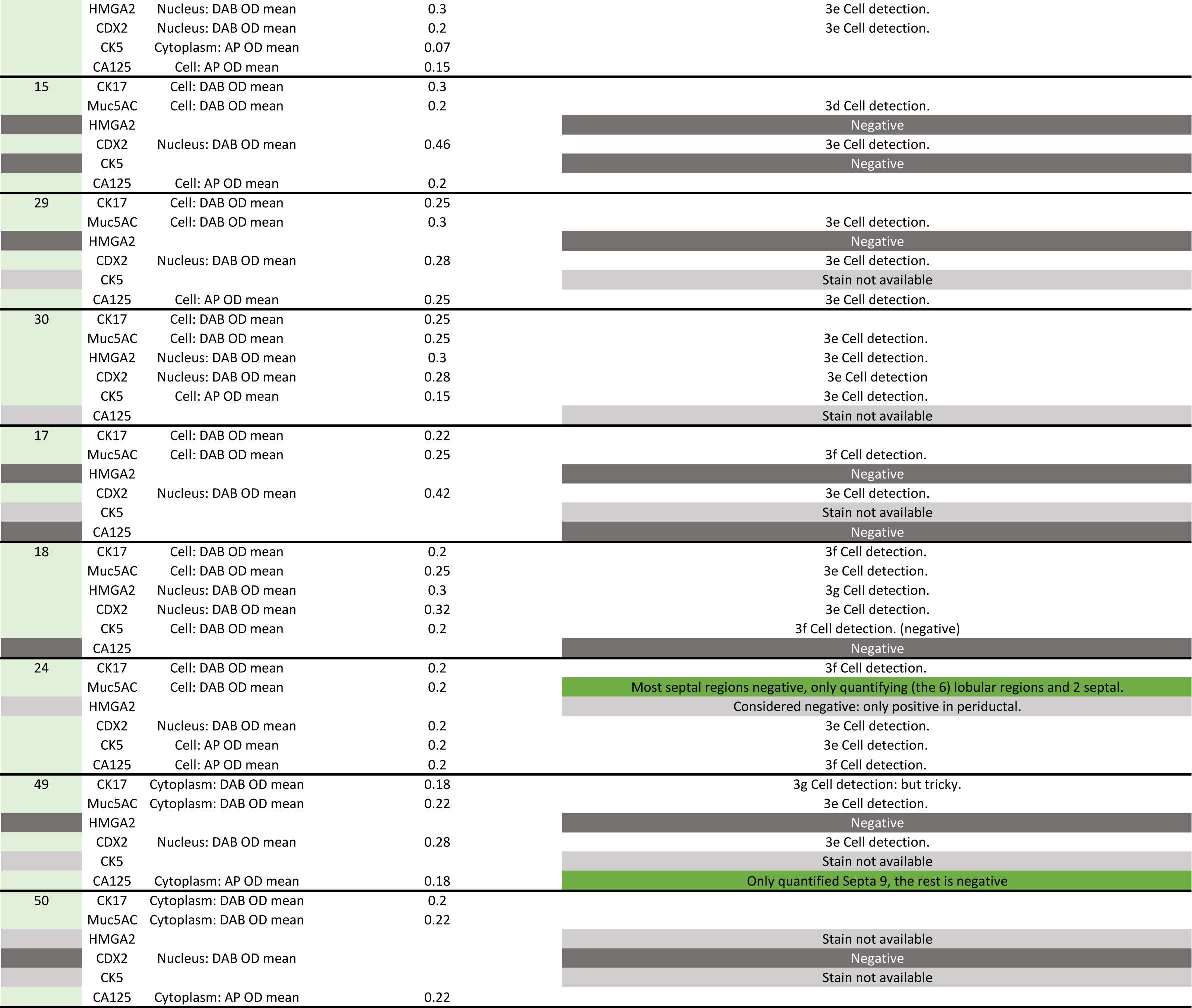
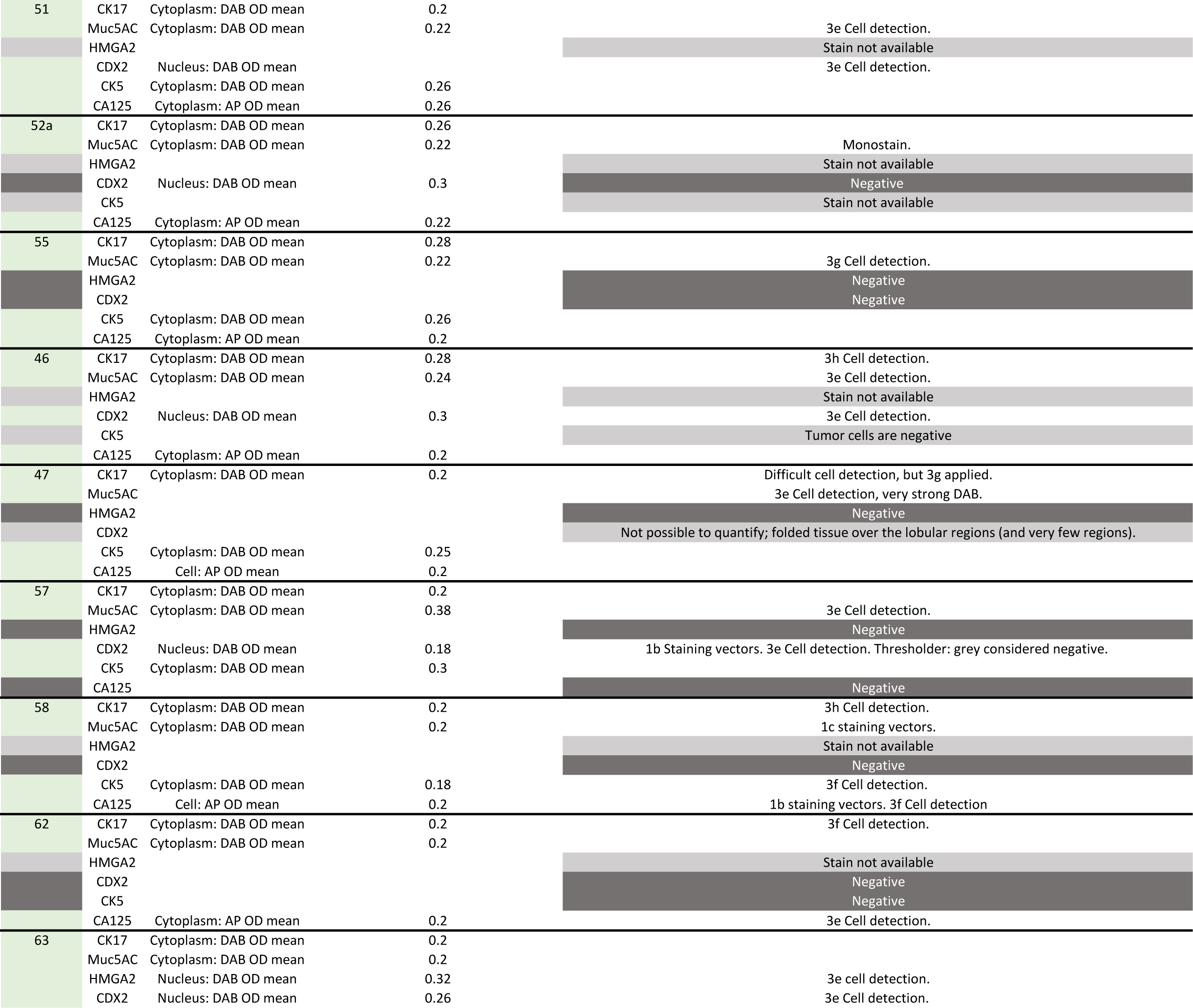
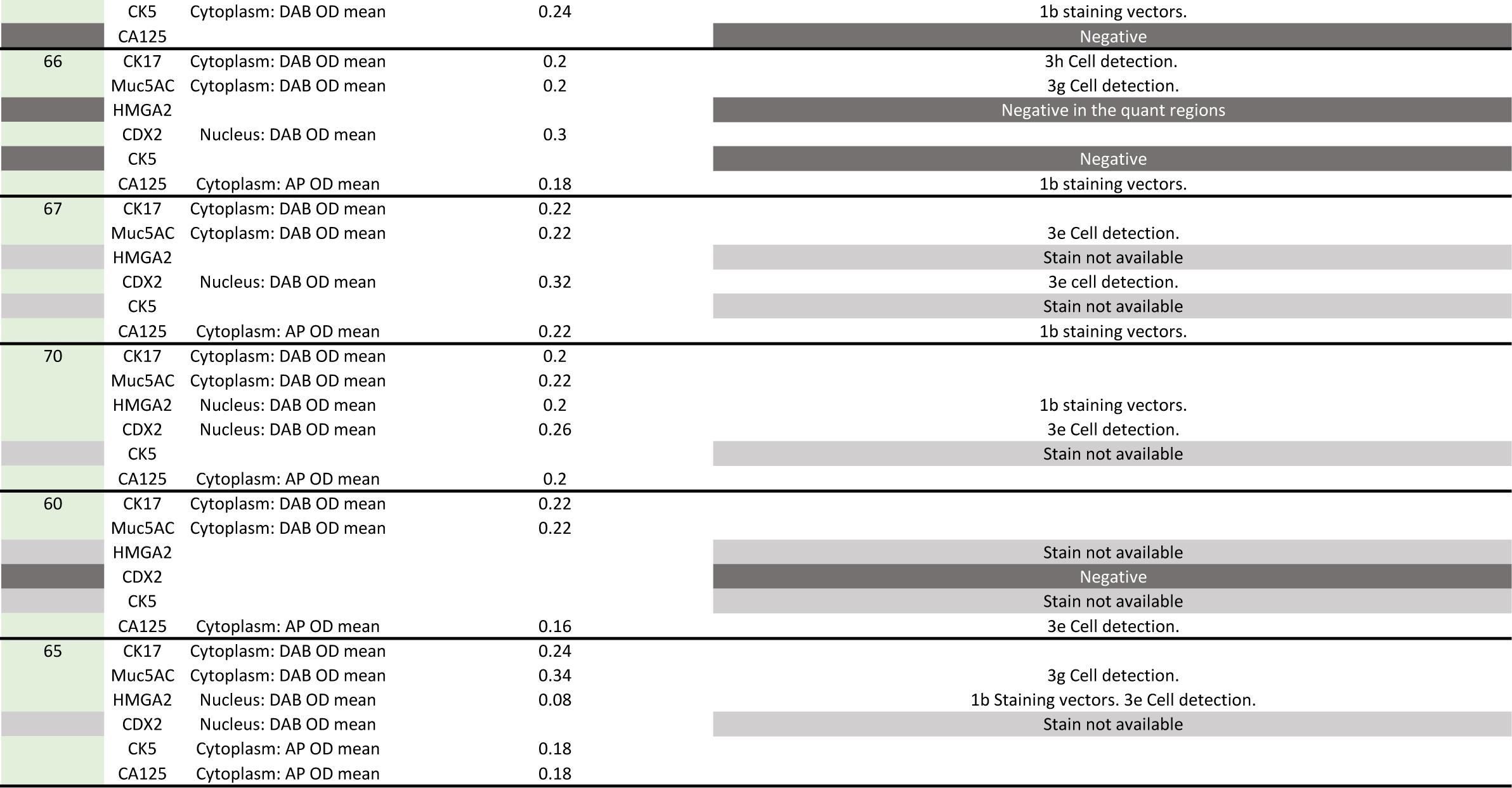
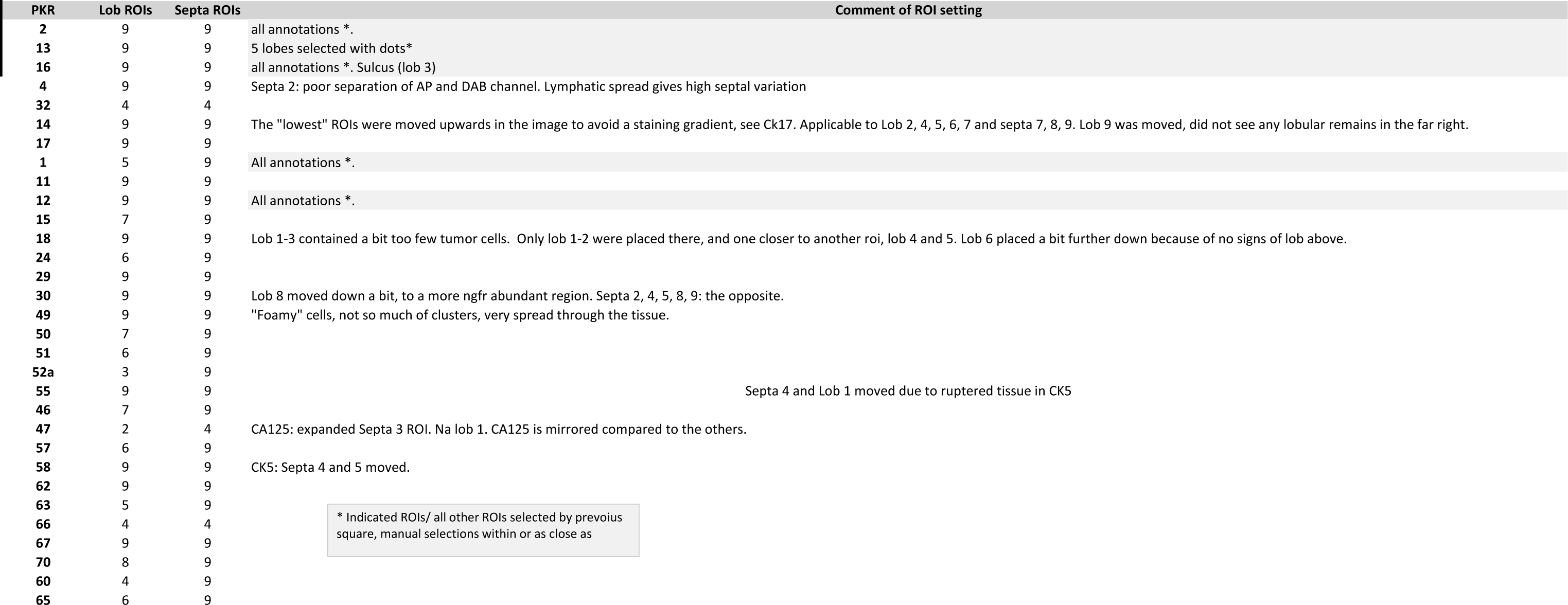

**Supplementary Table 2.** Convolutional neural network, generated with Aiforia Create v 5.5, model details, hyperparameters and error metrics.

|  |  |  |
| --- | --- | --- |
| Model objective |  | Recognition of main structures of pancreas |
| Ground truth |  | Acinar cells, acinar cells undergoing ADM, ducts, fibroblasts, large vessels, small vessels, nerves, immune cells, islets of Langerhans, tumor cells |
| Type (semantic segmentation) |  | Region |
| Complexity |  | Extra complex |
| Field of view | | 80 $\mu\text{m}$ |
| Training regions |  | 646 |
| Iterations |  | 2395 out of 3000 |
| Training parameters | Weight decay | 0.0001 |
|  | Mini batch size | 0 |
|  | Mini batches per iteration | 20 |
|  | Iterations without progress | 100 |
|  | Initial learning rate | 1 |
| Image augmentation | Scale (min/max) | -1/1 |
|  | Aspect ratio | 1 |
|  | Maximum shear | 1 |
|  | Luminance (min/max) | -1/1 |
|  | Contrast (min/max) | -1/1.01 |
|  | Max with balance change | 1 |
|  | Noise | 0 |
| Total area error % (FP/FN) |  | 3.46 (0.15/0.20) |
| Precision % |  | 98.37 |
| Sensitivity % |  | 97.85 |
| F1 Score % |  | 98.11 |
| Area error % |  | 0.35 |

ADM: acinar-to-ductal-metaplasia

**Supplementary table 3.**
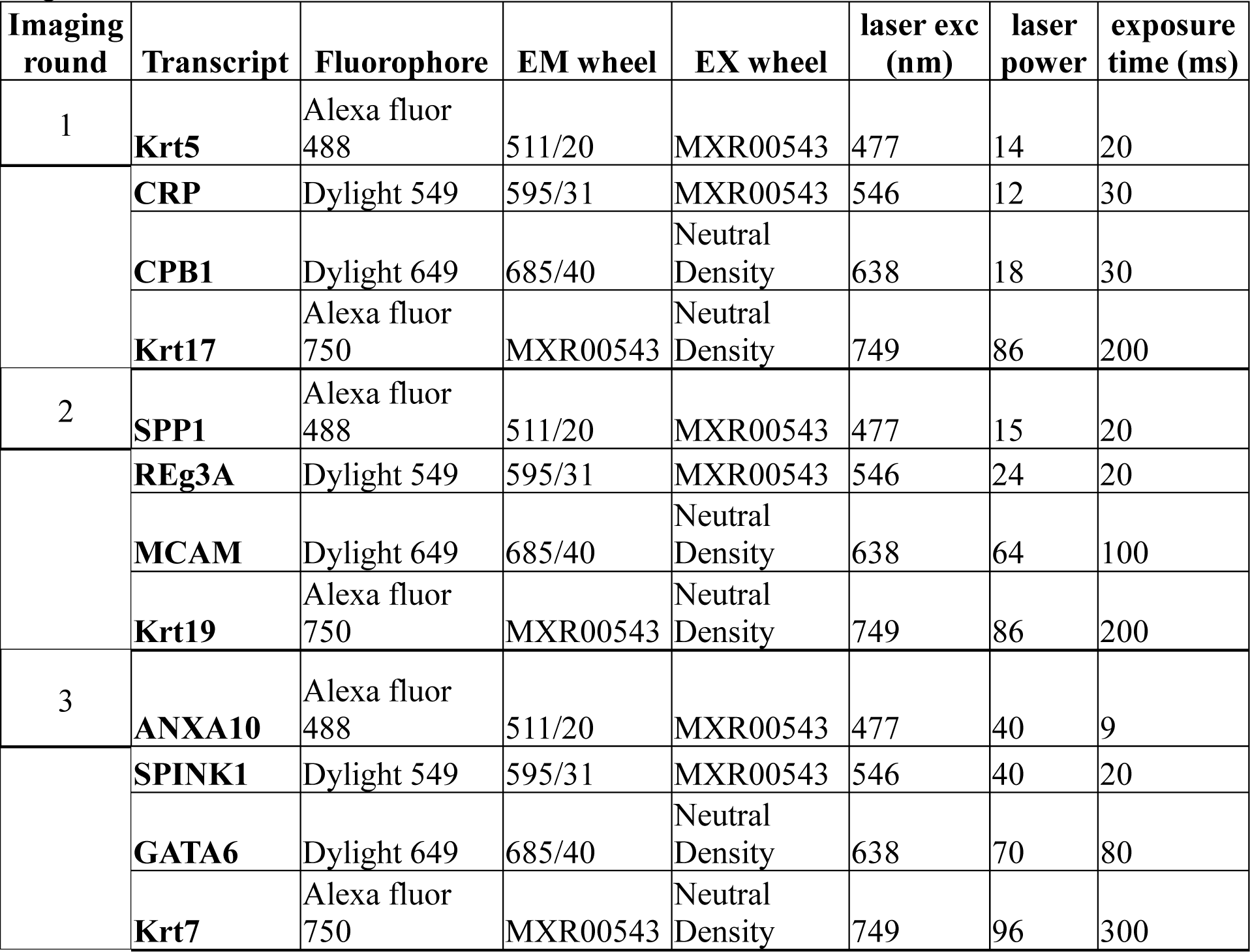
Image acquisition details for targets in the multiplex-RNA ISH. All fluorophore signals were acquired in widefield fluorescence modality with 1×1 binning at 12-bit depth. The DM wheel applied for all targets was a pentaband with the wavelength ranges: 441/30; 511/26; 593/37; 684/34 and 817/66.

| **Imaging round** | **Transcript** | **Fluorophore** | **EM wheel** | **EX wheel** | **laser exc (nm)** | **laser power** | **exposure time (ms)** |
| --- | --- | --- | --- | --- | --- | --- | --- |
| 1 | **Krt5** | Alexa fluor 488 | 511/20 | MXR00543 | 477 | 14 | 20 |
|  | **CRP** | Dylight 549 | 595/31 | MXR00543 | 546 | 12 | 30 |
|  | **CPB1** | Dylight 649 | 685/40 | Neutral Density | 638 | 18 | 30 |
|  | **Krt17** | Alexa fluor 750 | MXR00543 | Neutral Density | 749 | 86 | 200 |
| 2 | **SPP1** | Alexa fluor 488 | 511/20 | MXR00543 | 477 | 15 | 20 |
|  | **REg3A** | Dylight 549 | 595/31 | MXR00543 | 546 | 24 | 20 |
|  | **MCAM** | Dylight 649 | 685/40 | Neutral Density | 638 | 64 | 100 |
|  | **Krt19** | Alexa fluor 750 | MXR00543 | Neutral Density | 749 | 86 | 200 |
| 3 | **ANXA10** | Alexa fluor 488 | 511/20 | MXR00543 | 477 | 40 | 9 |
|  | **SPINK1** | Dylight 549 | 595/31 | MXR00543 | 546 | 40 | 20 |
|  | **GATA6** | Dylight 649 | 685/40 | Neutral Density | 638 | 70 | 80 |
|  | **Krt7** | Alexa fluor 750 | MXR00543 | Neutral Density | 749 | 96 | 300 |

**Supplementary table 4.**
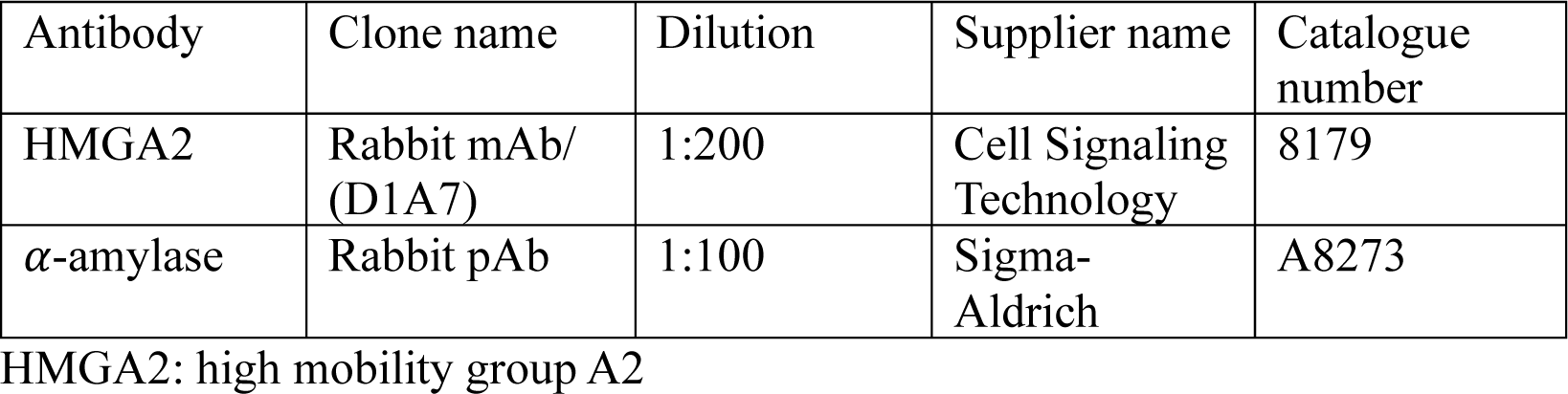
Details of the used primary antibodies in immunofluorescence of murine PDAC.

**Supplementary table 5.** Imaging details of targets from immunofluorescence of murine PDAC. All signals were acquired in widefield fluorescence modality with 1×1 binning at 12-bit depth. The DM wheel applied for all targets was a pentaband with the wavelengths ranges: 441/30; 511/26; 593/37; 684/34 and 817/66.

|  |  |  |
| --- | --- | --- |
| **Protein (antibody) target** | alpha-amylase | HMGA2 |
| **Secondary antibody/fluorophore** | (conj.) Alexa fluor 488 | Alexa fluor 647 |
| **EM wheel** | 511/20 | 685/40 |
| **EX wheel** | Neutal Density | MXR00543 (CELESTA-DA/FI/TR/Cy5/Cy7-A-OFF) |
| **Laser exc (nm)** | 477 | 638 |
| **Laser power** | 28 | 12 |
| **Exposure time (ms)** | 9 | 6 |

**Supplementary table 6.** Details of antibodies applied in the multiplex-immunofluorescence.

| **Antibody** | **Clone name** | **Dilution** | **Supplier name** | **Catalogue number** | **Lot number** |
| --- | --- | --- | --- | --- | --- |
| CD74 | Rabbit mAb/(D5N3 I) | 1:300 | Cell signaling | 77274T | Lot#1 |
| PDGFRα | Rabbit mAb(D13C 6) | 1:250 | Cell signaling | 5241S | Lot#4 |
| NGFR | Rabbit pAb | 1:500 | Atlas antibody | HPA004765 | Lot#000015823 |
| P53 | Mouse mAb(DO-7) | 1:300 | Leica biosystem | NCL-L-P53-D07 | Lot#P53-DO7-L-CE From 6065462 |
| ASMA | Mouse mAb(1A4) | 1:300 | Agilent | M0851 | Lot#20039209 |
| Vimentin | Rabbit mAb D21H3 | 1:500 | Cell signaling | 5741T | Lot#8 |

**Supplementary table 7.** The terminology for case ID:s between and example images (labelled T#_) the corresponding and plots (labelled PKR-_).

| Label on example image | Label in plots |
| --- | --- |
| T #1 | PKR-39 |
| T #2 | PKR-30 |
| T #3 | PKR-17 |
| T #4 | PKR-51 |
| T #5 | PKR-2 |
| T #6 | PKR-13 |
| T #7 | PKR-24 |
| T #8 | PKR-63 |
| T #9 | PKR-14 |
| T #10 | PKR-70 |
| T #11 | PKR-22 (only as example image) |
| T #12 | PKR-18 |

